# Codon Usage Bias Analysis of Human Papillomavirus 18’s L1 Protein and its Host Adaptability

**DOI:** 10.1101/2024.08.10.607454

**Authors:** Vinaya Shinde, Parminder Kaur, Swati Bankariya

## Abstract

Human Papillomavirus 18 (HPV 18) is known as a high-risk variant associated with cervical and anogenital malignancies. High-risk types HPV 18 and HPV 16 (human papillomavirus 16) play a major part in about 70 percent of cervical cancer worldwide (Ramakrishnan et al., 2015). The L1 protein of HPV 18 (HPV 18’s L1 protein), also known as major capsid L1 protein is targeted in the vaccine development against HPV 18 due to its non-oncogenic and non-infectious properties with self-assembly ability into virus-like particles. In the present analysis, an extensive codon usage bias analysis of HPV 18’s L1 protein and adaptation to its host human was conducted. The Effective number (Nc) Grand Average of Hydropathy (GRAVY), Index of Aromaticity (AROMO), and Codon Bias Index (CBI) values revealed no biases in codon usage of HPV 18’s L1 protein. The data of the Codon Adaptation Index (CAI), and Relative Codon Deoptimization Index (RCDI) indicate adaptation of HPV 18’s L1 protein according to its host human. The domination of selection pressure on codon usage of HPV 18’s L1 protein was demonstrated based on GC12 vs GC3, Nc vs GC3, and frequency of optimal codons (FOP). The Parity plot revealed that the genome of HPV 18’s L1 protein has a preference for purine over pyrimidine, that is G nucleotides over C, and no preference for A over T but A/T richness was observed in the genome of HPV 18’s L1 protein. In the Nucleotide composition, GC1 richness ultimately represents evolutionary aspects of codon usage. Furthermore, these findings can be used in currently ongoing vaccine development and gene therapy to design viral vectors.

## 1. Introduction

Human papillomavirus, also known as HPV, is an extremely oncogenic virus that causes most of the intraepithelial lesions as well as cervical cancer (Vinod Shinde et al., 2024). In the worldwide cervix, HPV 18 is thought to be responsible for 37% of cervical adenocarcinomas (ADC) and 12% of squamous cell carcinomas (SCC) (Chen et al., 2015). The HPV 18 is a double-stranded and circular-shaped DNA genome of size 7,857 base pairs (bp) with an approximate length of 8 kb (Graham, 2010). HPV 18 is a high-risk variant that has a significant role in the progression of cervical cancer, lung tumor tissue, and other cancers (“Human Papillomavirus Type 18 - an overview | ScienceDirect Topics,” n.d.). HPV 18 is one of the HPV variants that is involved in the progression of oral squamous cell carcinomas (OSCC) (Sri et al., 2021). HPV 18 can be transmitted through sexual activity and also through non-sexual activities like skin contact, mouth, fingers, and fomites (Petca et al., 2020).

The genome of HPV 18 encodes for almost 6 early gene regions ranging from E1 to E7, which are involved in the replication of the virus, whereas 2 late gene regions known as L1 and L2 are involved in the infectious process and assembly of HPV 18 as showed in **Fig.1**. The L1 protein of Human papillomavirus 18 (HPV 18’s L1 protein) also known as major capsid L1 protein, can self-assemble in the form of virus-like particles, also known as VLPs (Wang and Roden, 2013). The size of HPV 18’s L1 protein is ∼55 kD, which is involved in the initial interaction between the host human and HPV 18 (Buck et al., 2013). HPV 18’s L1 protein is important for viral entry and attachment, which is located at the late gene region of HPV 18’s life cycle to evade host immune surveillance (“L1 (Protein) - an overview | ScienceDirect Topics,” n.d.).

**Fig. 1.**
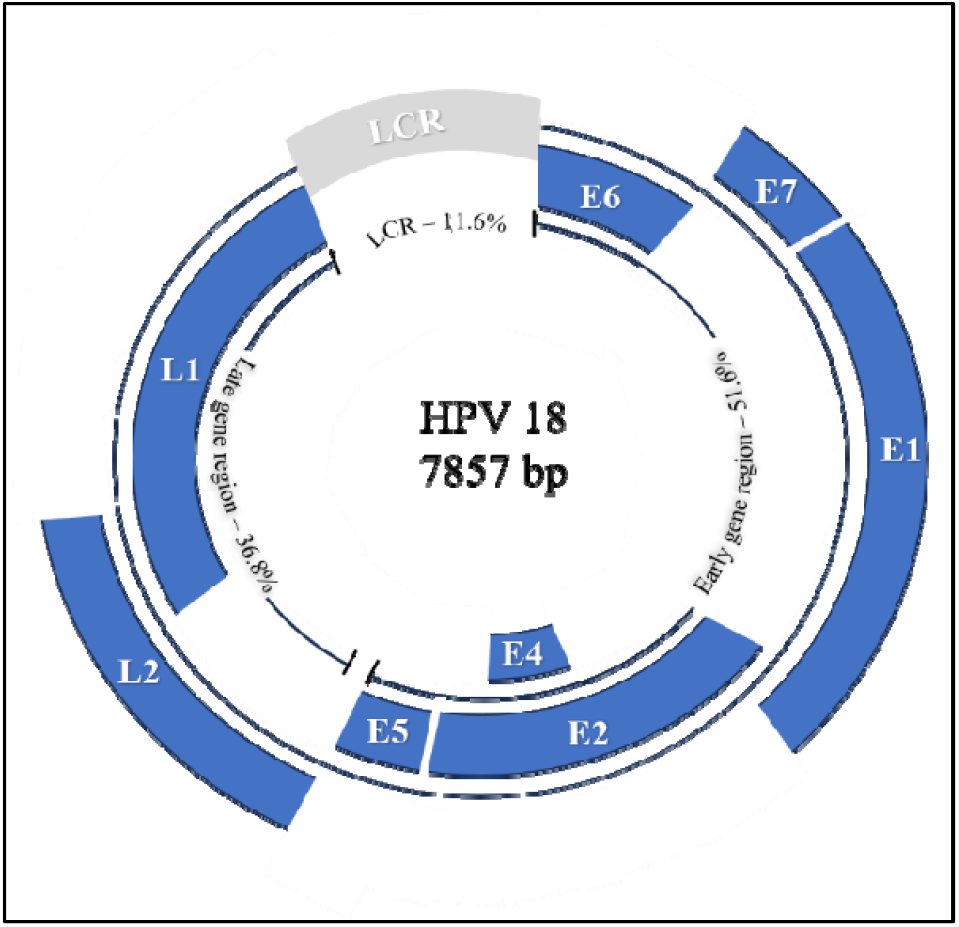
The viral circular genome structure of HPV 18 with 7857 bp. All viral ORFs are depicted in blue colour.

FDA-approved vaccines against HPV 18 infection are Cervarix (HPV Bivalent vaccine, Recombinant), Gardasil (HPV Quadrivalent vaccine, Recombinant), and Gardasil 9 (HPV 9 valent vaccine, Recomb nant) (U.S. Food and Drug Administration, 2018). This available vaccine against HPV 18 targets HPV 18’s L1 protein due to its non-oncogenic and immunogenic properties, HPV 18’s L1 protein can prevent HPV 18 infection by producing an antibody response (Ma et al., 2010).

The present analysis aimed to demonstrate the codon usage bias analysis of HPV 18’s L1 protein and adaptation to its host human, which was used to understand the biased codon usage, codon preferences, host adaptation, selection pressure, evolution, A/T richness, and preference for purine over pyrimidine in the genome of HPV 18’s L1 protein. Further, this analysis can support currently ongoing vaccine research against HPV 18 to target major capsid L1 protein and design optimized viral vectors in gene therapy. In the current study, we used isolated L1 protein in HPV 18 viral strains disseminated in the Netherlands at the beginning of vaccination to determine the baseline genetic variation in the Dutch population (King et al., 2016). This study on codon usage bias by HPV 18’s L1 protein aims to propose strategies to minimi e the virulence of HPV 18’s L1 protein and disclose its replicative efficacy and adaptation in host humans, which will facilitate the development of safe and effective vaccines. SAVE (Synthetic attenuated vaccine engineering) method uses the least preferred codons in vaccine production. This is the first publication to reveal the codon bias and host human adaptability of HPV 18’s L1 protein.

## 2. Materials and methods

### 2.1 Collection of Data

Overall all 108 Coding Sequences (CDSs) of HPV 18’s L1 protein were analyzed in this study. L1 protein Sequences of HPV 18 were derived from the data source, NCBI (National Center for Biotechnology Information) in the format of the FASTA file ranging from KU707718.1 to KU707825.1. The Codon Usage Table of the host human was derived from the Codon Usage Table in the CaiCAL server (Puigbò et al., 2008).

### 2.2 Nucleotide Content Analysis

Overall nucleotide content (A3, T3, C3, and G3, that is the individual nucleotides at the third location on the codon) of CDSs of HPV 18’s L1 protein was derived with the use of CodonW 1.4.2 program (Mareuil et al., 2017) (Khandia et al., 2019). GC1 (the nucleotide G, and C present at the first location on the codon), GC2 (the nucleotide G, and C present at the second location on the codon), GC3 (the nucleotide G, and C present at the third location on the codon) with GC12 (the average of GC1, and GC2) were retrieved with the use Bio.SeqUtils module in Bio-python. GC3s (third position in GC codon) was also derived from the CodonW 1.4.2 program.

### 2.3 Relative Synonymous Codon Usage Analysis (RSCU)

RSCU, which stands for Relative Synonymous Codon Usage, is a metric that analyzes and quantifies the codon bias in the synonymous codons inside genes or genomes. The concept is based on the fact that, in many organisms, specific synonymous codons coding for a particular amino acid are preferred over others (Khandia et al., 2019a). The frequency of amino acid or its length of the sequence doesn’t affect the RSCU value of CDSs. CodonW 1.4.2 program was used to derive the RSCU value of particular CDSs, the equation used to calculate the RSCU value is,

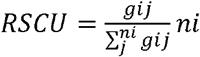

Where the observed number for the j^th^ amino acid’s i^th^ codon is g_ij_, which has synonymous codons of the ‘ni’ type. RSCU value above 1.6 reveals overrepresented codons while an RSCU value below 0.6 reveals underrepresented codons and RSCU output of codons in the range of 0.6 to 1.6 indicates randomly used or unbiasedness of codons.

### 2.4 Codon Adaptation Index (CAI)

The Codon Adaptation Index (CAI) is a statistic followed in molecular biology and bioinformatics to quantify the degree of adaptation of a codon usage inside a genome of the pathogen to the codon usage bias of a specific host. It determines how closely the codon usage bias of a given gene corresponds to the codon preferences reported in over-expressed genes inside the organism (He et al., 2021). A CAI value of 0 indicates low adaptation to the codon preferences identified in the reference genes, whereas a score of 1 indicates perfect alignment, signifying optimal adaptation. CAI scores were calculated with the use of CodonW 1.4.2. Program.

### 2.5 Frequency of optimal codons (FOP)

The frequency of optimal codons is the proportion of codons that are believed to be optimal for translation accuracy or correctness within a specific collection of genes or sequences and it was calculated with the use of CodonW 1.4.2. Program (Bahiri-Elitzur and Tuller, 2021). A value of 0 implies that none of the codons in the studied set are ideal, implying that codon usage is inefficient for translation efficiency and accuracy, which also indicates selection pressure. A value of 1 implies that all codons in the investigated set are optimal, implying that they are chosen primarily for translation accuracy or correctness based on criteria such as tRNA availability or ribosomal kinetics (Kunisawa et al., 1998).

### 2.6 Codon Bias Index (CBI)

The Codon Bias Index (CBI), which ranges from -1 to 1, evaluates the bias toward certain codons within a gene or genome. A CBI score of -1 shows no bias, implying equal use of all synonymous codons. As the CBI value is near 1, it represents a greater bias towards certain codons, with larger values suggesting a stronger preference for specific codons (Specht and Nissen, 1988). This codon bias might result from factors such as selection pressure, or gene expression levels. Codon 1.4.2. program was used to retrieve CBI values of HPV 18’s L1 protein.

### 2.7 Effective Number of Codon (Nc)

The Effective Number of Codons (Nc) is a study of statistical significance that determines the standard degree of codon use bias within a gene or genome. It quantifies the degree of bias toward specific codons, with lower Nc values indicating greater bias and higher values indicating more consistent codon usage (Wright, 1990). Nc value generally ranges from 20 to 61 (Sun et al., 2013). A lower Nc value, closer to 20, implies a significant bias, with a small number of codons favored for encoding every amino acid. The high Nc values, closer to 61, imply less bias and a more even distribution of codon usage between synonymous codons. NC values for all CDSs were derived from CodonW 1.4.2. Program, but it can be calculated manually by using the following formula,

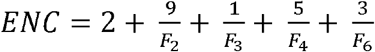

Where the values of Fk including 2, 3, 4, and 6 are indicated as amino acid degenerated K-fold pertained by Fk average value, F refers to the identical amino acid, and the probability of both randomly picked codons is studied to detect the identical amino acid (Bayer et al., 2012).

### 2.8 Aromaticity (AROMO) and Hydropathicity (GRAVY)

In codon usage analysis, the AROMO (Aromaticity Index) and GRAVY (Grand Average of Hydropathy) statistics provide useful information on the biochemical properties of proteins encoded by a gene or genome (Kumar et al., 2016). AROMO measures the abundance of aromatic amino acids on a parameter of the 0 to 1, with higher values revealing higher aromaticity, whereas GRAVY evaluates the hydropathic properties of proteins on a scale of -2 to +2, with positive values suggesting hydrophobicity and negative values suggesting hydrophilicity (Khandia et al., 2019a). These metrics help to understand protein structure, function, and cellular actions, with higher AROMO values indicating the presence of aromatic residues and positive GRAVY values suggesting the presence of hydrophobic inclinations, whereas negative GRAVY values suggesting the presence of hydrophilic inclinations. AROMO and GRAVY provide additional insight that is critical for understanding the physicochemical properties and genetic modifications of proteins across varied animals (Sheikh et al., 2020). All GRAVY and AROMO values of CDSs of HPV 18’s L1 protein were derived with the use of CodonW 1.4.2. Program.

### 2.9 Parity and Neutrality Analysis

Based on the ordinate, which is a bias of AT [A3/(A3 + T3)] and abscissa which is a bias of GC [G3/(G3 + C3)], a parity rule 2 (PR2) bias was plotted. Values of nucleotides A3, G3, T3, and C3 (A, G, T, and C nucleotides present at the third location on codon) were derived from CodonW1.4.2. Program and Matplotlib from Python were utilized to plot the Parity plot (Wang et al., 2022).

The effect on codon usage by translation selection and mutation bias has been studied using a neutrality plot. The contents of GC12 and GC3 were plotted along a regression line. The force of mutation is demonstrated by the direction of the regression line slope. Values of GC1, GC2, and GC3 and ultimately GC12 were derived from Bio.SeqUtils module in Bio Python and Microsoft Excel was used to plot the Neutrality plot (Khandia et al., 2019a).

## 3. Results and Discussion

### 3.1 Analysis of Nucleotide content of HPV 18’s L1 protein

Nucleotide composition affects the patterns in codon usage (Jenkins and Holmes, 2003). The nucleotide composition of 108 CDSs from HPV 18’s L1 protein was analyzed to study the effect on codon usage pattern by nucleotide composition. The general composition of nucleotide (GC1, GC2, GC3, T3, C3, A3, and G3) was analyzed in this analysis and compared in **Fig.2**. The appearance of nucleotide A3, G3, T3, and C3 at the third location on codon with average values was (0.29 ± 0.002), (0.51 ± 0.003), (0.21 ± 0.003) and (0.16 ± 0.002) respectively. One of the major components in codon usage analysis is GC content, codon usage patterns are found to be directly reflected by GC3 (He et al., 2016). The appearance of GC, which is GC1, GC2, and GC3 was observed with average values was (48.67 ± 0.08), (42.91 ± 0.09) and (32.02 ± 0.41) respectively. Higher GC nucleotide at the first position of codon represents the use of GC-rich codons in HPV 18’s L1 protein, which represents selection pressure on codon usage and ultimately represents the evolutionary aspects of codon usage.

**Fig. 2.**
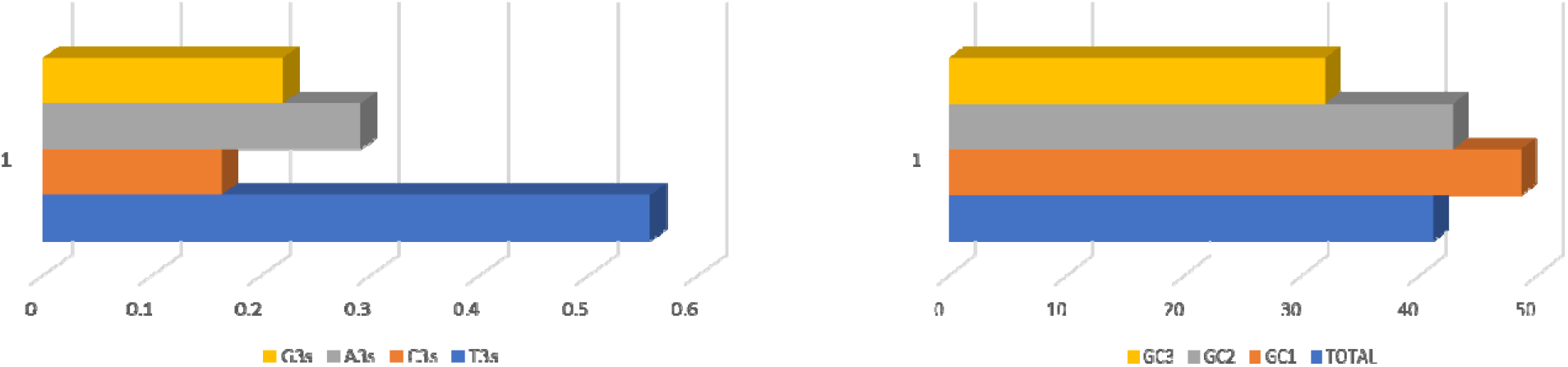
Overall nucleotide composition of A3, G3, T3, & C3 and GC1, GC2, GC3 & it’s total

### 3.2 A/T Richness in the genome of HPV 18’s L1 protein

The genome of HPV 18’s L1 protein was observed rich in A/T nucleotides when compared to C/G nucleotides. The mean values of A3, G3, T3, and C3 nucleotides were (0.29 ± 0.002), (0.51 ± 0.003), (0.21 ± 0.003), and (0.16 ± 0.002) respectively, representing the higher value of A and T nucleotides, when compared to C and G nucleotides. Python was used to analyze average of all CDSs of HPV 18’s L1 protein and standard deviation value of A, T, G, and C and it was recorded (440 ± 1.47), (562 ± 1.63), (366 ± 1.77), and (337 ± 1.17) respectively. Comparison of A/T with G/C represents the A/T-rich genome of HPV 18’s L1 protein.

### 3.3 Relative Synonymous codon usage (RSCU)

Relative Synonymous Codon Usage (RSCU) is a statistic that measures the bias or preference towards particular synonymous codons within HPV 18’s L1 protein sequences. It compares the analyzed frequency of each codon to the expected frequency, if all synonymous codons for a particular amino acid were used similarly (Wang et al., 2023). Overrepresented values of the RCSU are represented by light red color while underrepresented values of the RSCU are represented by yellow color.

Variation in RSCU values of CDSs of HPV 18’s L1 protein was visualized by plotting a heat map and all variation can be observed in the **Fig. 3**. Heat map reveals that, even inside the same gene preferences for codon show variation (Khandia et al., 2019b). Lower RSCU values are represented in dark blue colour while higher values of RSCU are represented in dark red colour. The main purpose of Heat maps is to visualize better the content or volume inside the data set or database with the use of colour code. The heat map was used to get a more reliable structure to calculate values, with a simple and general view of data.

**Fig. 3.**
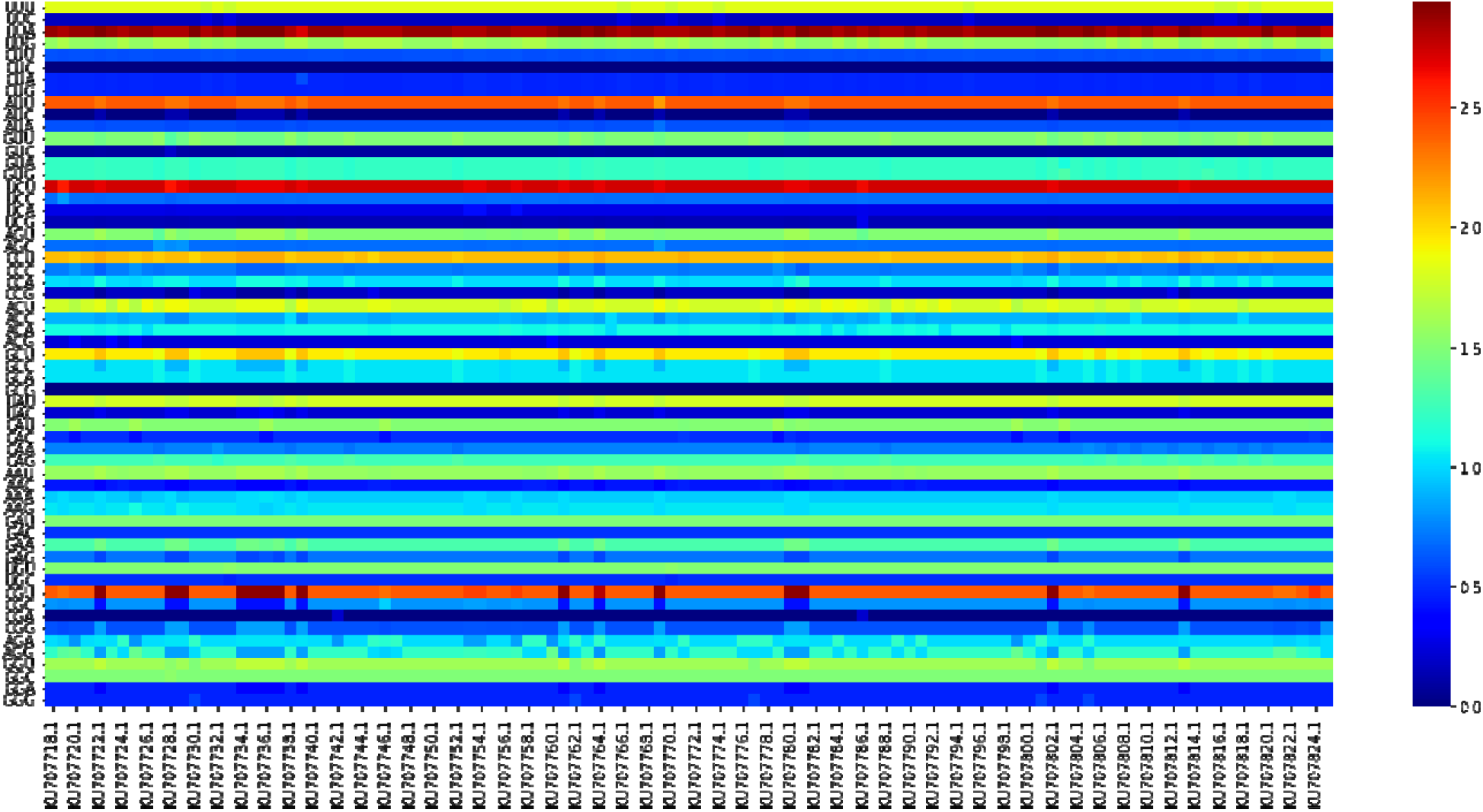
Visual representation of RSCU values of HPV 18’s L1 protein with the heatmap. Higher values are represented in dark red colour while all lower values are represented in dark blue colour.

### 3.4 No biases in codon usage of HPV 18’s L1 protein

Intra-genic codon bias was determined by calculating Nc values of HPV 18’s L1 protein genome. The highest Nc value and lowest Nc value in the present analysis were 44.77 and 43.14 respectively, while the mean value of Nc was (44.30 ± 0.43). Nc values almost close to 61 and more than 35 indicate no bias in the codon preferences of HPV 18’s L1 protein.

In the CBI analysis, The CBI score 1 reveals a substantial bias or favor for particular synonymous codons inside a given variety of genes or sequences (Choudhury et al., 2017). This implies that specific codons coding the same amino acid are utilized far more often than others, indicating selection force or optimization for the preferred codons. A CBI score of -1, on the other hand, implies that there is no bias or equal usage of synonymous codons in the investigated set of genes or sequences (Choudhury et al., 2018). This means that all codons coding the same amino acid are employed at similar frequencies, with no clear preference for any particular codon. The CBI value of (−0.080 ± 0.003) indicates an insignificant bias against specific synonymous codons within the investigated number of sequences. While the negative score suggests there is no severe bias toward specific codons, the modest divergence from zero implies that some codons are utilized less frequently than others. This could be due to slight evolutionary or functional influences impacting codon usage, although it is less evident than a positive CBI value, which indicates no significant bias toward certain codons.

The total average of all Grand Average of Hydropathy (GRAVY) values was (−0.28 ± 0.005), This negative GRAVY score indicates that, on average, proteins from viruses are slightly hydrophilic. Codons that encode hydrophilic amino acids, such as serine, asparagine, and threonine, may be favored in protein areas that are exposed to solvents or interact with other molecules. This preference may result in biased codon usage patterns that favor codons encoding hydrophilic amino acids.

The AROMO (Aromaticity Index) value of (0.105 ± 0.0004) indicates that synonymous codons in the investigated protein are used less often than usual. This study suggests a possible divergence from standard codon usage patterns, which could have repercussions for translation efficacy and protein expression. Further research into individual codon preferences and their effects on protein structure and function would shed light on the biological importance of this finding. The AROMO value is a useful statistic for determining codon use biases and their impact on protein structure and cell functions.

### 3.5 Study of HPV 18’s L1 protein adaptation

Virus adaptation according to the codon usage of the host and expression level of virus protein is evaluated with the use of CAI (Pan et al., 2020). Virus adaptation according to the codon usage of the host is considered higher when the CAI value is higher (approximately close to 1). The mean of CAI was (0.22 ± 0.002), which represents the low adaptation rate of HPV 18’s L1 protein according to its host human.

Comparison of the codon usage of HPV 18’s L1 protein and host human represents the effect on the expression of the gene or genome by affecting the codon bias of HPV 18’s L1 protein with (1.68 ± 0.02) value of RCDI. Observed co-evolution between host and virus can be studied with the evaluation of RCDI values. The higher value of RCDI in HPV 18’s L1 protein indicates a high adaptation of HPV 18’s L1 protein to their host human and also predicts the high translation rate of HPV 18’s L1 protein. The higher value of RCDI in HPV 18’s L1 protein concludes the similarity of codon usage of HPV 18’s L1 protein and the host human. Infectivity will be increased with an increase in adaptation and vice versa.

### 3.6 HPV 18’s L1 protein has selection pressure on codon usage

A study of mutational pressure on codon usage of HPV 18’s L1 protein was conducted with the comparison of GC12 and GC3 in the neutrality plot. The last nucleotide is changed and the translated protein remains the same in the synonymous codons. Mutational pressure can be studied with the fact that alteration in a nucleotide at the third codon location doesn’t affect amino acid alterations but amino acid alterations can be possible with alteration in nucleotide which is called selection pressure (Khandia et al., 2019a). The observed slope of the regression line from the GC12, and GC3 comparison was 0.0643 in **Fig. 4**, which indicated 6.43% mutational pressure (relative neutrality) on the HPV 18’s L1 protein. The mutational pressure of HPV 18’s L1 protein indicates that the selection pressure (relative constraint) of HPV 18’s L1 protein is 93.57%. In HPV 18’s L1 protein, the domination of selection pressure over the mutational pressure was observed.

**Fig. 4.**
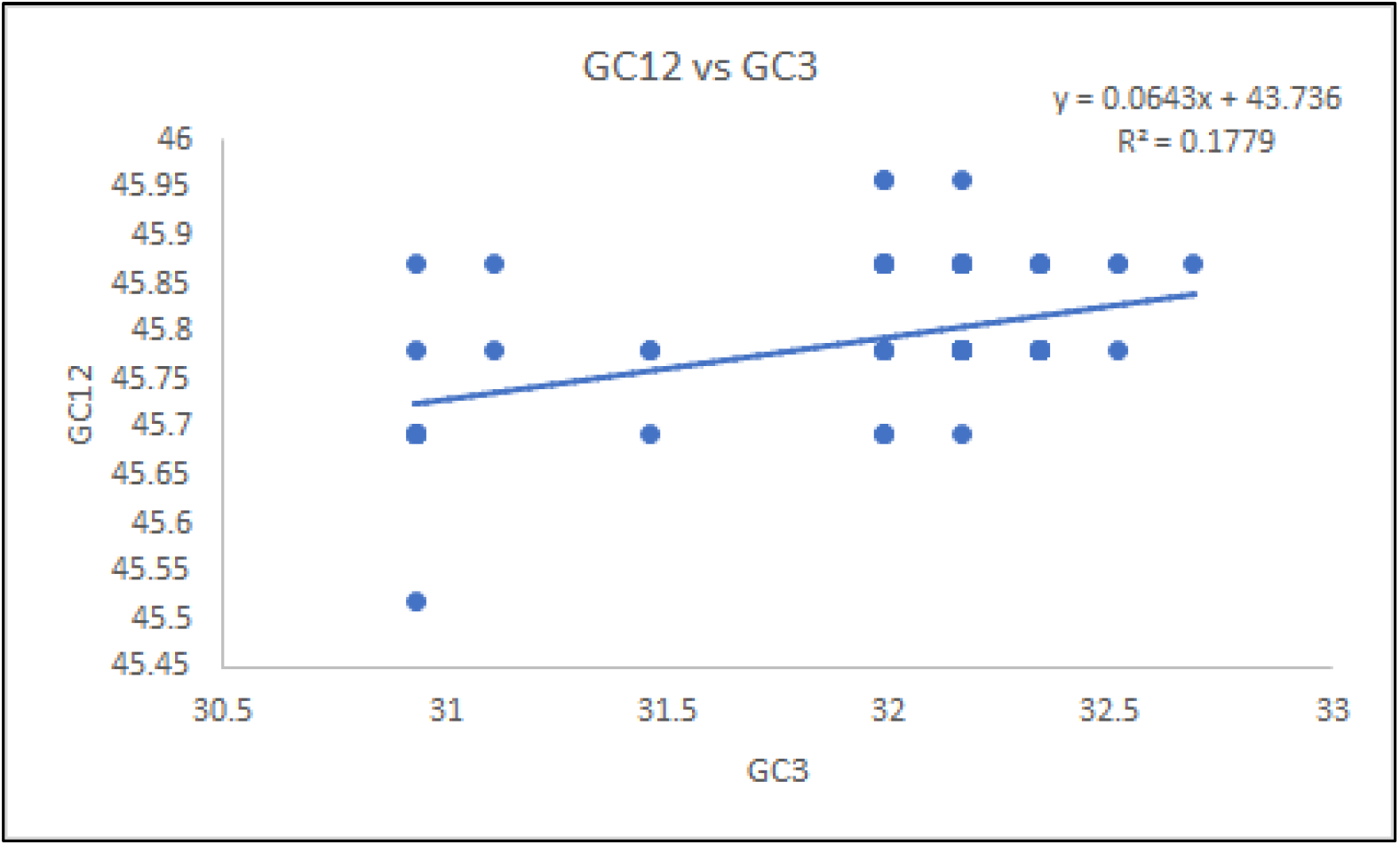
Neutrality plot analysis of L1 protein of HPV 18 by comparing GC12 and GC3 values.

A study of mutational and selection pressure on codon usage of HPV 18’s L1 protein was also studied with the comparison of Nc and GC3 plots. In the Nc and GC3 plot, the mutational and selection pressure on codon usage are concluded based on the data point position. Data points that lay above the expected curve suggest mutational pressure, whereas those that lay below reflect selection pressure on codon usage bias. In the Nc and GC3 analysis, all data points lay below the expected curve as shown in **Fig. 5**, which shows that the domination of selection pressure over the mutational pressure was observed.

**Fig. 5.**
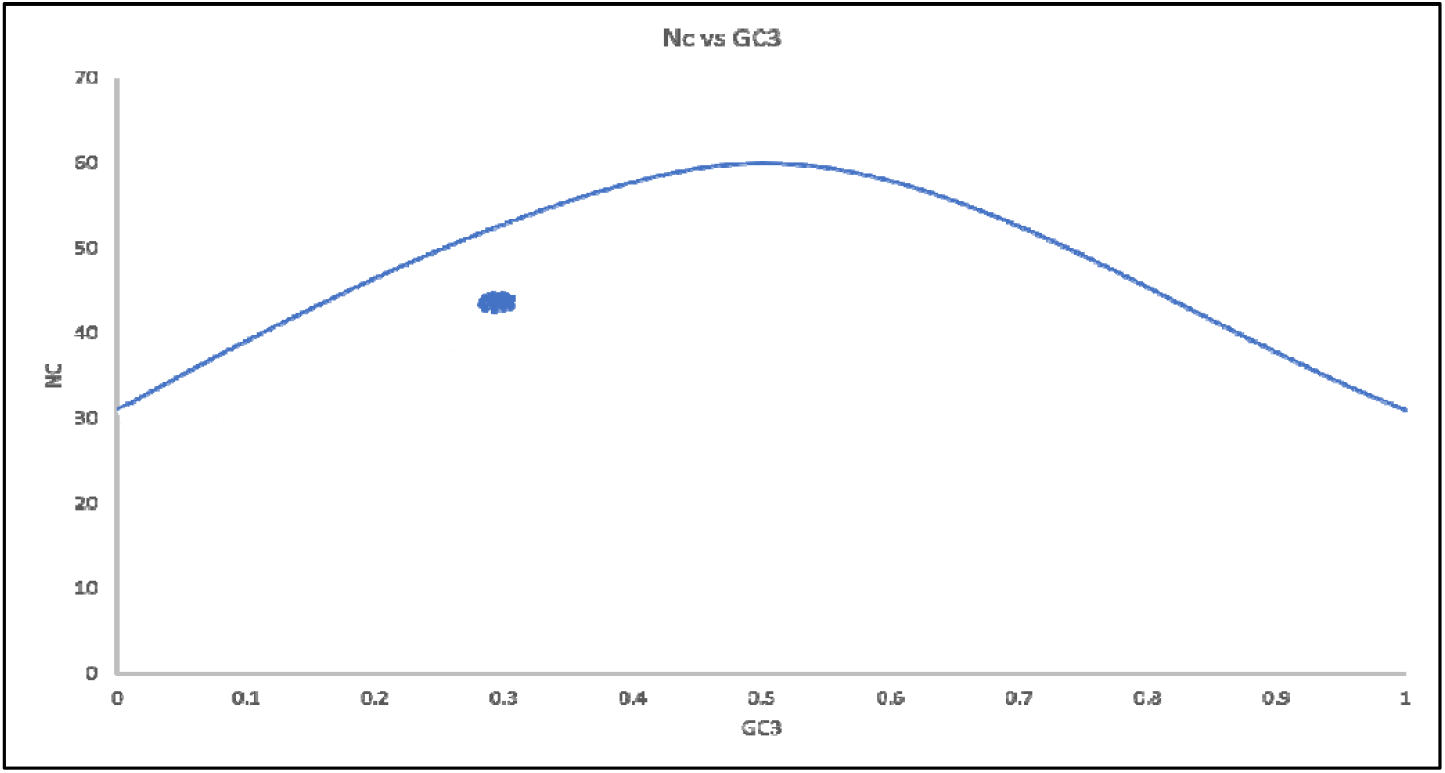
Nc vs GC3 plot analysis of L1 protein of HPV 18.

The frequency of optimal codons (FOP) represents the proportion of codons in a gene or genome that is considered to be optimal based on variables such as translation accuracy, effectiveness, or other selective pressures (10048480). Optimal codons are those that are preferred for reasons such as tRNA abundance, ribosomal kinetics, or other evolutionary or functional limitations. The frequency of optimum codons is frequently determined by comparing observed and expected codon usage based on tRNA frequencies or other relevant factors. The average frequency of the HPV 18’s L1 protein was (0.365 ± 0.002), indicating that around 36.5% of the codons in the examined set are regarded optimal for translation accuracy, efficacy, or other functional constraints. This implies that there is some selective pressure favoring the use of some codons over others in the investigated genes or sequences. In this sense, selective pressure refers to the influence of evolutionary factors or functional limitations on codon selection within a genome or gene set. The average FOP score of (0.365 ± 0.002) suggests that there is modest selective pressure favoring the use of optimum codons.

### 3.7 High preference for guanine (G) over cytosine (C) in HPV 18’s L1 protein

The Chargaff’s second parity rule (PR2) suggests that, the complementary nucleotides appear at almost equal rates in single-stranded DNA (Khandia et al., 2023). A parity plot was plotted against AT [A3/(A3 + T3)], and GC [G3/(G3 + C3)] to analyze preferences for purine and pyrimidine in the genome of HPV 18’s L1 protein.

Coordinates of both AT [A3/(A3 + T3)], and GC [G3/(G3 + C3)] intersect each other at the center of the plot with coordinate value 0.5 for both of them, where complementary nucleotides are the same (nucleotide A = T and G = C). The intersecting point at the coordinate value 0.5 also indicate no bias in the selection. The preference for purine over pyrimidine can concluded based on a bias value higher than 0.5, while a bias value lower than 0.5 represents a low preference for purine over pyrimidine. In the present plot, the average value of GC [G3/(G3 + C3)] was 0.58, which determines the preference for purine over pyrimidine and the average value of [A3/(A3 + T3)] was 0.35 as shown in **Fig. 6**, which determined the low preference for purine over pyrimidine in the HPV 18’s L1 protein. Hence, there will be a preference for G over C and no preference for A over C in the HPV 18’s L1 protein.

**Fig. 6.**
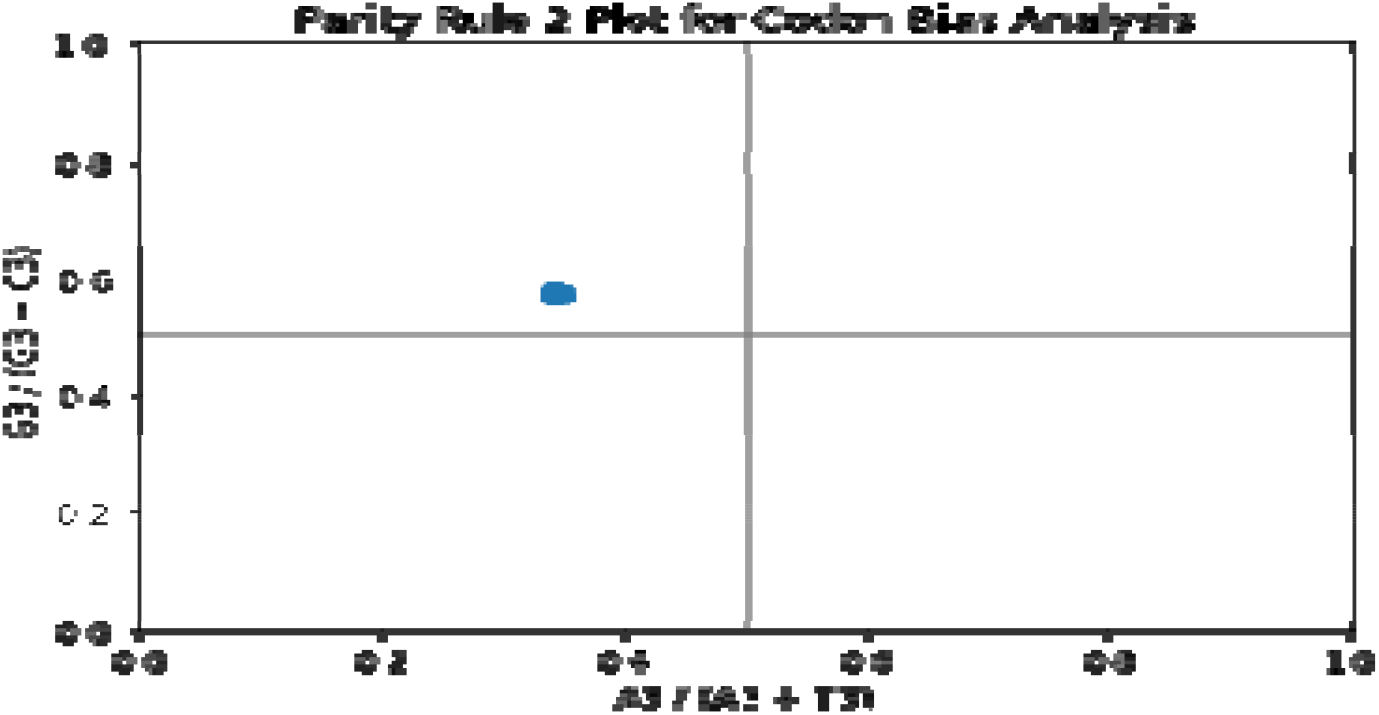
Parity plot analysis of L1 protein of HPV 18.

## 4. Conclusion

In this study, we investigated the codon usage of the L1 protein of Human Papillomavirus 18 (HPV 18’s L1 protein) to understand more about how this viral protein interacts with the host environment. Our investigation of multiple codon usage analyses, including Nc, CBI, GRAVY, and AROMO, indicated evidence of no bias in the codon usage pattern within the HPV 18 L1 protein genome. The abundance of aromatic and hydrophilic amino acids in HPV 18’s L1 proteins can drastically impact their efficiency, structural stability, and interactions with host cells, eventually influencing the virus’s infectiousness, pathogenicity, and immune evasion strategies. Furthermore, our analysis of the CAI and RCDI revealed a high level of adaptation of HPV 18’s L1 protein to its host environment. This adaptability most likely plays an important part in the ability of the virus to adapt and survive within its host.

Furthermore, the study we conducted on selection pressures on codon usage, as shown in GC12 vs GC3, Nc vs GC3, and FOP, revealed new insights into the evolutionary dynamics that shape the codon usage patterns of HPV 18’s L1 protein. The nucleotide composition analysis revealed an AT-rich genome of the HPV 18 L1 protein. Parity plot analysis indicated the preference for G over C (purine over pyrimidine) in the HPV 18’s L1 protein. In addition, the research revealed no significant preference for the A over C. Overall, this study improves the understanding of the evolution and host adaptability of HPV 18’s L1 protein, giving useful insights that may aid in the development of novel therapeutic techniques and vaccines for Human Papillomavirus infections.

## Supporting information

https://data.mendeley.com/preview/9ds7g93p7n?a=6206ea4a-fe06-49fc-8d61-32e38efe9804

## Contribution of authors

Vinaya Shinde collected, evaluated, and arranged all of the research data, as well as authored the publishing article. Dr. Parminder Kaur: Contributed considerable knowledge and meticulous evaluation of the data, analysis, and text, which helped to refine the research. Swati Bankariya demonstrated proper referencing throughout the research article’s preparation. The combined efforts of all authors were vital to the successful completion of the project.

## Supplementary Material

All supplemental material supporting this publication can be accessed through the following link : vindo, vinaya (2024), “Codon Usage Bias Analysis of Human Papillomavirus 18’s L1 Protein and its Host Adaptability”, Mendeley Data, V2, doi: 10.17632/9ds7g93p7n.2

## Acknowledgments

We sincerely thank Dr. Parminder Kaur for her essential contributions to the present study. Dr. Kaur contributed considerable knowledge and meticulously evaluated the data, analysis, and text, so playing a critical role in bringing the research to the highest possible standard.

