## Supplementary material for "Codon Usage Bias Analysis of Human Papillomavirus 18’s L1 Protein and its Host Adaptability": https://data.mendeley.com/preview/9ds7g93p7n?a=6206ea4a-fe06-49fc-8d61-32e38efe9804

**Supplementary table 1:** Nucleotides composition of L1 protein of Human Papillomavirus 18 genome at third codon position (A3s, T3s, G3s, and C3s) and GC1, GC2, and GC3 (Nucleotide present GC present at the first, second, and third position of codon) and GC3s (Nucleotide present at the third position of codon). CAI (Codon Adaptation Index), CBI (Codon Bias Index), FOP (Frequency of optimal codons), Nc (Effective Number), GRAVY and AROMO values of L1 protein of Human Papillomavirus 18 genome. G3/(A3+T3), A3/(A3+T3), and RCDI values of L1 protein of Human Papillomavirus 18 genome.

|  | **CDSs** | **GC1** | **GC2** | **GC3** | **GC12** | **T3s** | **C3s** | **A3s** | **G3s** | **CAI** | **CBI** | **Fop** | **Nc** | **GC3s** | **Gravy** | **AROMO** | **G3/(G3+C3)** | **A3/(A3+T3)** | **RCDI** |
| --- | --- | --- | --- | --- | --- | --- | --- | --- | --- | --- | --- | --- | --- | --- | --- | --- | --- | --- | --- |
| 1 | **KU707718.1** | 48.68 | 42.88 | 32.16 | 45.78 | 0.56 | 0.16 | 0.29 | 0.22 | 0.22 | -0.08 | 0.37 | 44.4 | 0.30 | -0.285 | 0.106 | 0.573 | 0.343 | 1.68 |
| 2 | **KU707719.1** | 48.68 | 43.06 | 32.34 | 45.87 | 0.55 | 0.17 | 0.29 | 0.22 | 0.22 | -0.08 | 0.36 | 44.7 | 0.30 | -0.282 | 0.106 | 0.573 | 0.344 | 1.68 |
| 3 | **KU707720.1** | 48.68 | 43.06 | 32.51 | 45.87 | 0.55 | 0.16 | 0.29 | 0.23 | 0.22 | -0.09 | 0.36 | 44.5 | 0.30 | -0.282 | 0.106 | 0.578 | 0.342 | 1.68 |
| 4 | **KU707721.1** | 48.68 | 42.88 | 32.16 | 45.78 | 0.56 | 0.16 | 0.29 | 0.22 | 0.22 | -0.08 | 0.37 | 44.4 | 0.30 | -0.285 | 0.106 | 0.573 | 0.343 | 1.68 |
| 5 | **KU707722.1** | 48.51 | 42.88 | 30.93 | 45.69 | 0.56 | 0.16 | 0.30 | 0.21 | 0.23 | -0.07 | 0.37 | 43.2 | 0.29 | -0.273 | 0.106 | 0.573 | 0.344 | 1.75 |
| 6 | **KU707723.1** | 48.68 | 42.88 | 32.34 | 45.78 | 0.55 | 0.16 | 0.29 | 0.22 | 0.22 | -0.09 | 0.36 | 44.6 | 0.30 | -0.285 | 0.106 | 0.575 | 0.344 | 1.68 |
| 7 | **KU707724.1** | 48.68 | 43.06 | 31.99 | 45.87 | 0.56 | 0.16 | 0.29 | 0.22 | 0.23 | -0.08 | 0.37 | 44.4 | 0.30 | -0.276 | 0.106 | 0.572 | 0.344 | 1.68 |
| 8 | **KU707725.1** | 48.68 | 43.06 | 32.69 | 45.87 | 0.55 | 0.16 | 0.29 | 0.23 | 0.22 | -0.09 | 0.36 | 44.5 | 0.30 | -0.282 | 0.106 | 0.580 | 0.340 | 1.68 |
| 9 | **KU707726.1** | 48.68 | 42.88 | 32.16 | 45.78 | 0.56 | 0.16 | 0.29 | 0.22 | 0.23 | -0.08 | 0.37 | 44.3 | 0.30 | -0.285 | 0.106 | 0.573 | 0.340 | 1.69 |
| 10 | **KU707727.1** | 48.68 | 42.88 | 32.34 | 45.78 | 0.55 | 0.17 | 0.29 | 0.22 | 0.22 | -0.08 | 0.37 | 44.6 | 0.30 | -0.274 | 0.106 | 0.573 | 0.344 | 1.68 |
| 11 | **KU707728.1** | 48.51 | 42.88 | 31.46 | 45.69 | 0.56 | 0.16 | 0.30 | 0.21 | 0.23 | -0.08 | 0.37 | 43.7 | 0.29 | -0.276 | 0.106 | 0.569 | 0.347 | 1.71 |
| 12 | **KU707729.1** | 48.68 | 42.88 | 31.46 | 45.78 | 0.56 | 0.16 | 0.29 | 0.22 | 0.23 | -0.08 | 0.37 | 43.5 | 0.29 | -0.275 | 0.106 | 0.575 | 0.344 | 1.72 |
| 13 | **KU707730.1** | 48.68 | 42.71 | 32.16 | 45.69 | 0.55 | 0.16 | 0.29 | 0.22 | 0.22 | -0.08 | 0.36 | 44.7 | 0.30 | -0.277 | 0.106 | 0.575 | 0.349 | 1.69 |
| 14 | **KU707731.1** | 48.68 | 42.88 | 32.34 | 45.78 | 0.56 | 0.17 | 0.29 | 0.22 | 0.23 | -0.08 | 0.37 | 44.7 | 0.30 | -0.286 | 0.107 | 0.570 | 0.342 | 1.67 |
| 15 | **KU707732.1** | 48.68 | 42.88 | 31.99 | 45.78 | 0.56 | 0.16 | 0.29 | 0.22 | 0.22 | -0.09 | 0.36 | 44.5 | 0.30 | -0.285 | 0.106 | 0.570 | 0.345 | 1.69 |
| 16 | **KU707733.1** | 48.68 | 43.06 | 32.34 | 45.87 | 0.56 | 0.17 | 0.29 | 0.22 | 0.23 | -0.08 | 0.37 | 44.6 | 0.30 | -0.280 | 0.106 | 0.570 | 0.342 | 1.67 |
| 17 | **KU707734.1** | 48.51 | 42.88 | 30.93 | 45.69 | 0.56 | 0.16 | 0.30 | 0.21 | 0.23 | -0.07 | 0.37 | 43.2 | 0.29 | -0.273 | 0.106 | 0.573 | 0.344 | 1.75 |
| 18 | **KU707735.1** | 48.51 | 42.88 | 30.93 | 45.69 | 0.56 | 0.16 | 0.30 | 0.21 | 0.23 | -0.07 | 0.37 | 43.2 | 0.29 | -0.273 | 0.106 | 0.573 | 0.344 | 1.75 |
| 19 | **KU707736.1** | 48.15 | 42.88 | 30.93 | 45.52 | 0.56 | 0.16 | 0.30 | 0.21 | 0.23 | -0.07 | 0.37 | 43.2 | 0.29 | -0.271 | 0.107 | 0.573 | 0.344 | 1.75 |
| 20 | **KU707737.1** | 48.51 | 42.88 | 30.93 | 45.69 | 0.56 | 0.16 | 0.30 | 0.21 | 0.23 | -0.07 | 0.37 | 43.2 | 0.29 | -0.273 | 0.106 | 0.573 | 0.344 | 1.75 |
| 21 | **KU707738.1** | 48.68 | 42.88 | 32.34 | 45.78 | 0.55 | 0.17 | 0.29 | 0.22 | 0.22 | -0.08 | 0.36 | 44.6 | 0.30 | -0.274 | 0.106 | 0.573 | 0.344 | 1.68 |
| 22 | **KU707739.1** | 48.86 | 42.88 | 31.11 | 45.87 | 0.56 | 0.16 | 0.29 | 0.21 | 0.23 | -0.07 | 0.37 | 43.5 | 0.29 | -0.275 | 0.106 | 0.576 | 0.342 | 1.72 |
| 23 | **KU707740.1** | 48.68 | 42.88 | 32.16 | 45.78 | 0.56 | 0.16 | 0.29 | 0.22 | 0.22 | -0.08 | 0.37 | 44.4 | 0.30 | -0.285 | 0.106 | 0.573 | 0.343 | 1.68 |
| 24 | **KU707741.1** | 48.68 | 42.88 | 32.16 | 45.78 | 0.56 | 0.16 | 0.29 | 0.22 | 0.22 | -0.08 | 0.37 | 44.4 | 0.30 | -0.285 | 0.106 | 0.573 | 0.343 | 1.68 |
| 25 | **KU707742.1** | 48.86 | 42.88 | 32.16 | 45.87 | 0.56 | 0.16 | 0.29 | 0.22 | 0.22 | -0.08 | 0.37 | 44.5 | 0.30 | -0.285 | 0.106 | 0.573 | 0.343 | 1.68 |
| 26 | **KU707743.1** | 48.68 | 42.88 | 32.16 | 45.78 | 0.55 | 0.16 | 0.29 | 0.22 | 0.22 | -0.08 | 0.36 | 44.5 | 0.30 | -0.274 | 0.106 | 0.576 | 0.344 | 1.68 |
| 27 | **KU707744.1** | 48.68 | 42.88 | 32.16 | 45.78 | 0.56 | 0.16 | 0.29 | 0.22 | 0.22 | -0.08 | 0.36 | 44.4 | 0.30 | -0.281 | 0.106 | 0.575 | 0.343 | 1.68 |
| 28 | **KU707745.1** | 48.68 | 43.06 | 32.16 | 45.87 | 0.55 | 0.16 | 0.29 | 0.22 | 0.23 | -0.08 | 0.37 | 44.6 | 0.30 | -0.276 | 0.106 | 0.575 | 0.345 | 1.68 |
| 29 | **KU707746.1** | 48.68 | 43.23 | 31.99 | 45.96 | 0.56 | 0.16 | 0.29 | 0.22 | 0.22 | -0.08 | 0.37 | 44.3 | 0.30 | -0.279 | 0.106 | 0.571 | 0.344 | 1.68 |
| 30 | **KU707747.1** | 48.68 | 42.88 | 31.99 | 45.78 | 0.56 | 0.16 | 0.29 | 0.22 | 0.22 | -0.08 | 0.36 | 44.4 | 0.30 | -0.281 | 0.106 | 0.573 | 0.345 | 1.68 |
| 31 | **KU707748.1** | 48.68 | 42.88 | 32.16 | 45.78 | 0.56 | 0.16 | 0.29 | 0.22 | 0.22 | -0.08 | 0.37 | 44.4 | 0.30 | -0.285 | 0.106 | 0.573 | 0.343 | 1.68 |
| 32 | **KU707749.1** | 48.68 | 42.88 | 32.16 | 45.78 | 0.56 | 0.16 | 0.29 | 0.22 | 0.22 | -0.08 | 0.37 | 44.4 | 0.30 | -0.285 | 0.106 | 0.573 | 0.343 | 1.68 |
| 33 | **KU707750.1** | 48.68 | 42.88 | 32.16 | 45.78 | 0.56 | 0.16 | 0.29 | 0.22 | 0.22 | -0.08 | 0.37 | 44.4 | 0.30 | -0.285 | 0.106 | 0.573 | 0.343 | 1.68 |
| 34 | **KU707751.1** | 48.68 | 42.88 | 32.16 | 45.78 | 0.56 | 0.16 | 0.29 | 0.22 | 0.22 | -0.08 | 0.37 | 44.4 | 0.30 | -0.285 | 0.106 | 0.573 | 0.343 | 1.68 |
| 35 | **KU707752.1** | 48.68 | 42.88 | 32.16 | 45.78 | 0.55 | 0.16 | 0.29 | 0.22 | 0.22 | -0.08 | 0.36 | 44.5 | 0.30 | -0.274 | 0.106 | 0.576 | 0.344 | 1.68 |
| 36 | **KU707753.1** | 48.68 | 42.88 | 32.16 | 45.78 | 0.56 | 0.16 | 0.29 | 0.22 | 0.23 | -0.08 | 0.37 | 44.6 | 0.30 | -0.292 | 0.106 | 0.572 | 0.343 | 1.67 |
| 37 | **KU707754.1** | 48.68 | 42.88 | 32.16 | 45.78 | 0.56 | 0.16 | 0.29 | 0.22 | 0.23 | -0.08 | 0.37 | 44.6 | 0.30 | -0.292 | 0.106 | 0.572 | 0.343 | 1.67 |
| 38 | **KU707755.1** | 48.68 | 42.88 | 32.16 | 45.78 | 0.56 | 0.16 | 0.29 | 0.22 | 0.22 | -0.08 | 0.37 | 44.4 | 0.30 | -0.285 | 0.106 | 0.573 | 0.343 | 1.68 |
| 39 | **KU707756.1** | 48.86 | 42.88 | 32.16 | 45.87 | 0.56 | 0.16 | 0.29 | 0.22 | 0.22 | -0.08 | 0.37 | 44.4 | 0.30 | -0.280 | 0.106 | 0.573 | 0.343 | 1.68 |
| 40 | **KU707757.1** | 48.68 | 42.88 | 32.16 | 45.78 | 0.56 | 0.16 | 0.29 | 0.22 | 0.23 | -0.08 | 0.37 | 44.6 | 0.30 | -0.292 | 0.106 | 0.572 | 0.343 | 1.67 |
| 41 | **KU707758.1** | 48.68 | 43.06 | 31.99 | 45.87 | 0.56 | 0.16 | 0.29 | 0.22 | 0.23 | -0.08 | 0.37 | 44.4 | 0.30 | -0.276 | 0.106 | 0.572 | 0.344 | 1.68 |
| 42 | **KU707759.1** | 48.68 | 43.06 | 31.99 | 45.87 | 0.56 | 0.16 | 0.29 | 0.22 | 0.23 | -0.08 | 0.37 | 44.4 | 0.30 | -0.276 | 0.106 | 0.572 | 0.344 | 1.68 |
| 43 | **KU707760.1** | 48.68 | 42.88 | 32.51 | 45.78 | 0.55 | 0.16 | 0.29 | 0.23 | 0.22 | -0.09 | 0.36 | 44.6 | 0.30 | -0.285 | 0.106 | 0.578 | 0.342 | 1.75 |
| 44 | **KU707761.1** | 48.51 | 42.88 | 30.93 | 45.69 | 0.56 | 0.16 | 0.30 | 0.21 | 0.23 | -0.07 | 0.37 | 43.2 | 0.29 | -0.273 | 0.106 | 0.573 | 0.344 | 1.68 |
| 45 | **KU707762.1** | 48.68 | 42.71 | 31.99 | 45.69 | 0.55 | 0.16 | 0.29 | 0.22 | 0.22 | -0.09 | 0.36 | 44.5 | 0.30 | -0.277 | 0.106 | 0.573 | 0.348 | 1.68 |
| 46 | **KU707763.1** | 48.68 | 42.88 | 32.16 | 45.78 | 0.56 | 0.16 | 0.29 | 0.22 | 0.22 | -0.08 | 0.37 | 44.4 | 0.30 | -0.285 | 0.106 | 0.573 | 0.343 | 1.69 |
| 47 | **KU707764.1** | 48.51 | 43.06 | 30.93 | 45.78 | 0.56 | 0.16 | 0.30 | 0.21 | 0.23 | -0.07 | 0.37 | 43.2 | 0.29 | -0.274 | 0.106 | 0.573 | 0.345 | 1.68 |
| 48 | **KU707765.1** | 48.68 | 42.88 | 32.34 | 45.78 | 0.56 | 0.17 | 0.29 | 0.22 | 0.23 | -0.08 | 0.37 | 44.4 | 0.30 | -0.285 | 0.106 | 0.569 | 0.341 | 1.75 |
| 49 | **KU707766.1** | 48.68 | 42.88 | 32.34 | 45.78 | 0.56 | 0.17 | 0.29 | 0.22 | 0.23 | -0.08 | 0.37 | 44.7 | 0.30 | -0.286 | 0.107 | 0.570 | 0.342 | 1.67 |
| 50 | **KU707767.1** | 48.68 | 42.88 | 32.16 | 45.78 | 0.56 | 0.16 | 0.29 | 0.22 | 0.22 | -0.08 | 0.37 | 44.4 | 0.30 | -0.285 | 0.106 | 0.573 | 0.343 | 1.68 |
| 51 | **KU707768.1** | 48.68 | 42.88 | 32.16 | 45.78 | 0.56 | 0.16 | 0.29 | 0.22 | 0.22 | -0.08 | 0.37 | 44.4 | 0.30 | -0.285 | 0.106 | 0.573 | 0.343 | 1.68 |
| 52 | **KU707769.1** | 48.68 | 42.88 | 31.46 | 45.78 | 0.56 | 0.16 | 0.30 | 0.22 | 0.23 | -0.08 | 0.37 | 43.8 | 0.29 | -0.275 | 0.106 | 0.575 | 0.347 | 1.72 |
| 53 | **KU707770.1** | 49.03 | 42.88 | 32.16 | 45.96 | 0.56 | 0.16 | 0.29 | 0.22 | 0.22 | -0.08 | 0.36 | 44.6 | 0.30 | -0.282 | 0.106 | 0.575 | 0.343 | 1.67 |
| 54 | **KU707771.1** | 48.68 | 43.06 | 32.34 | 45.87 | 0.56 | 0.17 | 0.29 | 0.22 | 0.23 | -0.08 | 0.37 | 44.6 | 0.30 | -0.280 | 0.106 | 0.570 | 0.342 | 1.67 |
| 55 | **KU707772.1** | 48.68 | 43.23 | 31.99 | 45.96 | 0.56 | 0.16 | 0.29 | 0.22 | 0.22 | -0.08 | 0.37 | 44.4 | 0.30 | -0.273 | 0.106 | 0.571 | 0.344 | 1.68 |
| 56 | **KU707773.1** | 48.68 | 42.88 | 32.16 | 45.78 | 0.56 | 0.16 | 0.29 | 0.22 | 0.22 | -0.08 | 0.37 | 44.4 | 0.30 | -0.285 | 0.106 | 0.573 | 0.343 | 1.68 |
| 57 | **KU707774.1** | 48.68 | 42.88 | 32.16 | 45.78 | 0.56 | 0.16 | 0.29 | 0.22 | 0.22 | -0.08 | 0.37 | 44.4 | 0.30 | -0.285 | 0.106 | 0.573 | 0.343 | 1.68 |
| 58 | **KU707775.1** | 48.68 | 42.88 | 32.34 | 45.78 | 0.56 | 0.17 | 0.29 | 0.22 | 0.23 | -0.08 | 0.37 | 44.7 | 0.30 | -0.286 | 0.107 | 0.570 | 0.342 | 1.67 |
| 59 | **KU707776.1** | 48.68 | 42.88 | 32.16 | 45.78 | 0.56 | 0.16 | 0.29 | 0.22 | 0.22 | -0.08 | 0.37 | 44.4 | 0.30 | -0.285 | 0.106 | 0.573 | 0.343 | 1.68 |
| 60 | **KU707777.1** | 48.68 | 42.88 | 31.99 | 45.78 | 0.56 | 0.16 | 0.29 | 0.22 | 0.22 | -0.08 | 0.37 | 44.4 | 0.30 | -0.285 | 0.106 | 0.570 | 0.345 | 1.68 |
| 61 | **KU707778.1** | 48.68 | 42.71 | 31.99 | 45.69 | 0.55 | 0.16 | 0.29 | 0.22 | 0.22 | -0.09 | 0.36 | 44.5 | 0.30 | -0.277 | 0.106 | 0.573 | 0.348 | 1.69 |
| 62 | **KU707779.1** | 48.68 | 43.06 | 31.99 | 45.87 | 0.56 | 0.16 | 0.29 | 0.22 | 0.23 | -0.08 | 0.37 | 44.4 | 0.30 | -0.276 | 0.106 | 0.572 | 0.344 | 1.68 |
| 63 | **KU707780.1** | 48.68 | 43.06 | 32.16 | 45.87 | 0.56 | 0.16 | 0.29 | 0.22 | 0.22 | -0.08 | 0.36 | 44.3 | 0.30 | -0.282 | 0.106 | 0.572 | 0.343 | 1.69 |
| 64 | **KU707781.1** | 48.68 | 42.88 | 31.11 | 45.78 | 0.56 | 0.16 | 0.29 | 0.21 | 0.23 | -0.07 | 0.37 | 43.3 | 0.29 | -0.275 | 0.106 | 0.576 | 0.342 | 1.74 |
| 65 | **KU707782.1** | 48.68 | 43.06 | 30.93 | 45.87 | 0.56 | 0.16 | 0.30 | 0.21 | 0.23 | -0.07 | 0.37 | 43.1 | 0.29 | -0.269 | 0.106 | 0.573 | 0.345 | 1.75 |
| 66 | **KU707783.1** | 48.68 | 42.88 | 32.16 | 45.78 | 0.56 | 0.16 | 0.29 | 0.22 | 0.22 | -0.08 | 0.37 | 44.4 | 0.30 | -0.285 | 0.106 | 0.573 | 0.343 | 1.68 |
| 67 | **KU707784.1** | 48.68 | 42.88 | 32.16 | 45.78 | 0.56 | 0.16 | 0.29 | 0.22 | 0.23 | -0.08 | 0.37 | 44.3 | 0.30 | -0.285 | 0.106 | 0.573 | 0.340 | 1.69 |
| 68 | **KU707785.1** | 48.68 | 43.06 | 31.99 | 45.87 | 0.56 | 0.16 | 0.29 | 0.22 | 0.23 | -0.08 | 0.37 | 44.4 | 0.30 | -0.276 | 0.106 | 0.572 | 0.344 | 1.68 |
| 69 | **KU707786.1** | 48.68 | 42.88 | 32.16 | 45.78 | 0.56 | 0.16 | 0.29 | 0.22 | 0.23 | -0.08 | 0.37 | 44.3 | 0.30 | -0.285 | 0.106 | 0.573 | 0.340 | 1.69 |
| 70 | **KU707787.1** | 48.86 | 42.88 | 32.16 | 45.87 | 0.55 | 0.16 | 0.29 | 0.22 | 0.22 | -0.08 | 0.36 | 44.8 | 0.30 | -0.281 | 0.106 | 0.575 | 0.346 | 1.68 |
| 71 | **KU707788.1** | 48.68 | 42.88 | 32.16 | 45.78 | 0.56 | 0.16 | 0.29 | 0.22 | 0.22 | -0.08 | 0.37 | 44.4 | 0.30 | -0.285 | 0.106 | 0.573 | 0.343 | 1.68 |
| 72 | **KU707789.1** | 48.68 | 42.88 | 32.34 | 45.78 | 0.55 | 0.17 | 0.29 | 0.22 | 0.22 | -0.08 | 0.36 | 44.6 | 0.30 | -0.274 | 0.106 | 0.573 | 0.344 | 1.68 |
| 73 | **KU707790.1** | 48.68 | 43.06 | 31.99 | 45.87 | 0.56 | 0.16 | 0.29 | 0.22 | 0.23 | -0.08 | 0.37 | 44.4 | 0.30 | -0.276 | 0.106 | 0.572 | 0.344 | 1.68 |
| 74 | **KU707791.1** | 48.68 | 42.88 | 32.16 | 45.78 | 0.56 | 0.16 | 0.29 | 0.22 | 0.22 | -0.08 | 0.37 | 44.4 | 0.30 | -0.285 | 0.106 | 0.573 | 0.343 | 1.68 |
| 75 | **KU707792.1** | 48.68 | 43.06 | 31.99 | 45.87 | 0.56 | 0.16 | 0.29 | 0.22 | 0.23 | -0.08 | 0.37 | 44.4 | 0.30 | -0.276 | 0.106 | 0.572 | 0.344 | 1.68 |
| 76 | **KU707793.1** | 48.68 | 42.88 | 32.16 | 45.78 | 0.56 | 0.16 | 0.29 | 0.22 | 0.22 | -0.08 | 0.37 | 44.4 | 0.30 | -0.285 | 0.106 | 0.573 | 0.343 | 1.68 |
| 77 | **KU707794.1** | 48.68 | 42.88 | 32.16 | 45.78 | 0.56 | 0.16 | 0.29 | 0.22 | 0.23 | -0.08 | 0.37 | 44.3 | 0.30 | -0.285 | 0.106 | 0.573 | 0.340 | 1.69 |
| 78 | **KU707795.1** | 48.68 | 42.88 | 32.16 | 45.78 | 0.56 | 0.16 | 0.29 | 0.22 | 0.22 | -0.08 | 0.37 | 44.4 | 0.30 | -0.285 | 0.106 | 0.573 | 0.343 | 1.68 |
| 79 | **KU707796.1** | 48.68 | 42.88 | 32.34 | 45.78 | 0.56 | 0.17 | 0.29 | 0.22 | 0.23 | -0.08 | 0.37 | 44.7 | 0.30 | -0.286 | 0.107 | 0.570 | 0.342 | 1.67 |
| 80 | **KU707797.1** | 48.68 | 42.88 | 32.16 | 45.78 | 0.56 | 0.16 | 0.29 | 0.22 | 0.22 | -0.08 | 0.37 | 44.4 | 0.30 | -0.285 | 0.106 | 0.573 | 0.343 | 1.68 |
| 81 | **KU707798.1** | 48.68 | 42.88 | 32.16 | 45.78 | 0.56 | 0.16 | 0.29 | 0.22 | 0.22 | -0.08 | 0.37 | 44.4 | 0.30 | -0.285 | 0.106 | 0.573 | 0.343 | 1.68 |
| 82 | **KU707799.1** | 48.68 | 42.88 | 32.16 | 45.78 | 0.56 | 0.16 | 0.29 | 0.22 | 0.23 | -0.08 | 0.37 | 44.3 | 0.30 | -0.285 | 0.106 | 0.573 | 0.340 | 1.69 |
| 83 | **KU707800.1** | 48.68 | 43.06 | 32.51 | 45.87 | 0.55 | 0.16 | 0.29 | 0.23 | 0.22 | -0.09 | 0.36 | 44.5 | 0.30 | -0.282 | 0.106 | 0.578 | 0.342 | 1.68 |
| 84 | **KU707801.1** | 48.68 | 42.88 | 32.16 | 45.78 | 0.56 | 0.16 | 0.29 | 0.22 | 0.22 | -0.08 | 0.37 | 44.4 | 0.30 | -0.285 | 0.106 | 0.573 | 0.343 | 1.68 |
| 85 | **KU707802.1** | 48.68 | 42.71 | 31.99 | 45.69 | 0.55 | 0.16 | 0.29 | 0.22 | 0.22 | -0.09 | 0.36 | 44.5 | 0.30 | -0.277 | 0.106 | 0.573 | 0.348 | 1.69 |
| 86 | **KU707803.1** | 48.51 | 42.88 | 30.93 | 45.69 | 0.56 | 0.16 | 0.30 | 0.21 | 0.23 | -0.07 | 0.37 | 43.2 | 0.29 | -0.273 | 0.106 | 0.573 | 0.344 | 1.75 |
| 87 | **KU707804.1** | 48.68 | 43.06 | 32.34 | 45.87 | 0.56 | 0.16 | 0.29 | 0.22 | 0.22 | -0.08 | 0.36 | 44.3 | 0.30 | -0.282 | 0.106 | 0.575 | 0.341 | 1.68 |
| 88 | **KU707805.1** | 48.68 | 42.88 | 32.16 | 45.78 | 0.56 | 0.16 | 0.29 | 0.22 | 0.22 | -0.08 | 0.37 | 44.4 | 0.30 | -0.285 | 0.106 | 0.573 | 0.343 | 1.68 |
| 89 | **KU707806.1** | 48.68 | 42.88 | 31.99 | 45.78 | 0.55 | 0.16 | 0.29 | 0.22 | 0.22 | -0.09 | 0.36 | 44.7 | 0.30 | -0.279 | 0.106 | 0.573 | 0.348 | 1.69 |
| 90 | **KU707807.1** | 48.86 | 42.88 | 32.16 | 45.87 | 0.56 | 0.16 | 0.29 | 0.22 | 0.22 | -0.08 | 0.37 | 44.4 | 0.30 | -0.280 | 0.106 | 0.573 | 0.343 | 1.68 |
| 91 | **KU707808.1** | 48.68 | 42.88 | 32.16 | 45.78 | 0.55 | 0.16 | 0.29 | 0.22 | 0.22 | -0.08 | 0.36 | 44.5 | 0.30 | -0.274 | 0.106 | 0.576 | 0.344 | 1.68 |
| 92 | **KU707809.1** | 48.68 | 42.88 | 32.16 | 45.78 | 0.56 | 0.16 | 0.29 | 0.22 | 0.22 | -0.08 | 0.37 | 44.4 | 0.30 | -0.285 | 0.106 | 0.573 | 0.343 | 1.68 |
| 93 | **KU707810.1** | 48.68 | 42.88 | 32.34 | 45.78 | 0.55 | 0.17 | 0.29 | 0.22 | 0.22 | -0.08 | 0.36 | 44.6 | 0.30 | -0.274 | 0.106 | 0.573 | 0.344 | 1.68 |
| 94 | **KU707811.1** | 48.68 | 42.88 | 32.16 | 45.78 | 0.56 | 0.16 | 0.29 | 0.22 | 0.22 | -0.08 | 0.37 | 44.4 | 0.30 | -0.285 | 0.106 | 0.573 | 0.343 | 1.68 |
| 95 | **KU707812.1** | 48.68 | 42.88 | 32.16 | 45.78 | 0.56 | 0.16 | 0.29 | 0.22 | 0.22 | -0.08 | 0.36 | 44.4 | 0.30 | -0.281 | 0.106 | 0.575 | 0.343 | 1.68 |
| 96 | **KU707813.1** | 48.68 | 43.06 | 32.34 | 45.87 | 0.55 | 0.16 | 0.29 | 0.23 | 0.22 | -0.08 | 0.37 | 44.5 | 0.30 | -0.278 | 0.106 | 0.579 | 0.342 | 1.68 |
| 97 | **KU707814.1** | 48.51 | 42.88 | 30.93 | 45.69 | 0.56 | 0.16 | 0.30 | 0.21 | 0.23 | -0.07 | 0.37 | 43.2 | 0.29 | -0.273 | 0.106 | 0.573 | 0.344 | 1.75 |
| 98 | **KU707815.1** | 48.68 | 42.88 | 32.16 | 45.78 | 0.56 | 0.16 | 0.29 | 0.22 | 0.22 | -0.08 | 0.37 | 44.4 | 0.30 | -0.285 | 0.106 | 0.573 | 0.343 | 1.68 |
| 99 | **KU707816.1** | 48.68 | 42.88 | 32.34 | 45.78 | 0.55 | 0.16 | 0.29 | 0.23 | 0.22 | -0.08 | 0.36 | 44.5 | 0.30 | -0.274 | 0.106 | 0.579 | 0.344 | 1.67 |
| 100 | **KU707817.1** | 48.68 | 42.88 | 32.34 | 45.78 | 0.56 | 0.17 | 0.29 | 0.22 | 0.23 | -0.08 | 0.37 | 44.7 | 0.30 | -0.286 | 0.107 | 0.570 | 0.342 | 1.68 |
| 101 | **KU707818.1** | 48.68 | 42.88 | 32.34 | 45.78 | 0.56 | 0.17 | 0.29 | 0.22 | 0.23 | -0.08 | 0.37 | 44.7 | 0.30 | -0.286 | 0.107 | 0.570 | 0.342 | 1.67 |
| 102 | **KU707819.1** | 48.68 | 42.88 | 32.34 | 45.78 | 0.55 | 0.17 | 0.29 | 0.22 | 0.22 | -0.08 | 0.36 | 44.6 | 0.30 | -0.274 | 0.106 | 0.573 | 0.344 | 1.68 |
| 103 | **KU707820.1** | 48.68 | 42.88 | 32.34 | 45.78 | 0.56 | 0.17 | 0.29 | 0.22 | 0.23 | -0.08 | 0.37 | 44.7 | 0.30 | -0.286 | 0.107 | 0.570 | 0.342 | 1.67 |
| 104 | **KU707821.1** | 48.68 | 42.88 | 31.99 | 45.78 | 0.55 | 0.16 | 0.29 | 0.22 | 0.22 | -0.08 | 0.36 | 44.4 | 0.30 | -0.274 | 0.106 | 0.573 | 0.346 | 1.68 |
| 105 | **KU707822.1** | 48.68 | 43.06 | 32.16 | 45.87 | 0.55 | 0.16 | 0.29 | 0.22 | 0.22 | -0.08 | 0.36 | 44.5 | 0.30 | -0.282 | 0.106 | 0.576 | 0.344 | 1.70 |
| 106 | **KU707823.1** | 48.68 | 43.06 | 32.16 | 45.87 | 0.55 | 0.16 | 0.29 | 0.22 | 0.22 | -0.08 | 0.36 | 44.5 | 0.30 | -0.282 | 0.106 | 0.576 | 0.344 | 1.68 |
| 107 | **KU707824.1** | 48.68 | 42.88 | 32.16 | 45.78 | 0.56 | 0.16 | 0.29 | 0.22 | 0.22 | -0.08 | 0.37 | 44.4 | 0.30 | -0.285 | 0.106 | 0.573 | 0.343 | 1.68 |
| 108 | **KU707825.1** | 48.68 | 43.06 | 32.16 | 45.87 | 0.56 | 0.16 | 0.29 | 0.22 | 0.23 | -0.08 | 0.37 | 44.3 | 0.30 | -0.287 | 0.106 | 0.572 | 0.343 | 1.69 |
