## Supplementary material for "Codon Usage Bias Analysis of Human Papillomavirus 18’s L1 Protein and its Host Adaptability": https://data.mendeley.com/preview/9ds7g93p7n?a=6206ea4a-fe06-49fc-8d61-32e38efe9804

**Supplementary table 2.1:** Values of Relative Synonymous Codon Usage (RSCU) for accession number from KU707718.1 to KU707730.1 of L1 protein of Human Papillomavirus 18.

| **Amino acids** | **Codons** | **KU707718.1** | **KU707719.1** | **KU707720.1** | **KU707721.1** | **KU707722.1** | **KU707723.1** | **KU707724.1** | **KU707725.1** | **KU707726.1** | **KU707727.1** | **KU707728.1** | **KU707729.1** | **KU707730.1** |
| --- | --- | --- | --- | --- | --- | --- | --- | --- | --- | --- | --- | --- | --- | --- |
| Phenylalanine(F) | UUU | 1.83 | 1.83 | 1.83 | 1.83 | 1.83 | 1.83 | 1.83 | 1.83 | 1.83 | 1.83 | 1.83 | 1.83 | 1.83 |
|  | UUC | 0.17 | 0.17 | 0.17 | 0.17 | 0.17 | 0.17 | 0.17 | 0.17 | 0.17 | 0.17 | 0.17 | 0.17 | 0.17 |
| Leucine(L) | UUA | 2.88 | 2.82 | 2.88 | 2.88 | 2.94 | 2.88 | 2.82 | 2.88 | 2.88 | 2.82 | 2.82 | 2.82 | 2.94 |
|  | UUG | 1.56 | 1.65 | 1.56 | 1.56 | 1.53 | 1.56 | 1.65 | 1.56 | 1.56 | 1.65 | 1.65 | 1.65 | 1.53 |
|  | CUU | 0.6 | 0.59 | 0.6 | 0.6 | 0.59 | 0.6 | 0.59 | 0.6 | 0.6 | 0.59 | 0.59 | 0.59 | 0.59 |
|  | CUC | 0 | 0 | 0 | 0 | 0 | 0 | 0 | 0 | 0 | 0 | 0 | 0 | 0 |
|  | CUA | 0.48 | 0.47 | 0.48 | 0.48 | 0.47 | 0.48 | 0.47 | 0.48 | 0.48 | 0.47 | 0.47 | 0.47 | 0.47 |
|  | CUG | 0.48 | 0.47 | 0.48 | 0.48 | 0.47 | 0.48 | 0.47 | 0.48 | 0.48 | 0.47 | 0.47 | 0.47 | 0.47 |
| Isoleucine(I) | AUU | 2.4 | 2.4 | 2.4 | 2.4 | 2.31 | 2.4 | 2.4 | 2.4 | 2.4 | 2.4 | 2.31 | 2.31 | 2.4 |
|  | AUC | 0 | 0 | 0 | 0 | 0.12 | 0 | 0 | 0 | 0 | 0 | 0.12 | 0.12 | 0 |
|  | AUA | 0.6 | 0.6 | 0.6 | 0.6 | 0.58 | 0.6 | 0.6 | 0.6 | 0.6 | 0.6 | 0.58 | 0.58 | 0.6 |
| Valine(V) | GUU | 1.49 | 1.49 | 1.49 | 1.49 | 1.43 | 1.49 | 1.49 | 1.49 | 1.49 | 1.55 | 1.33 | 1.43 | 1.55 |
|  | GUC | 0.09 | 0.09 | 0.09 | 0.09 | 0.1 | 0.09 | 0.09 | 0.09 | 0.09 | 0.09 | 0.19 | 0.1 | 0.09 |
|  | GUA | 1.21 | 1.21 | 1.21 | 1.21 | 1.24 | 1.21 | 1.21 | 1.21 | 1.21 | 1.18 | 1.24 | 1.24 | 1.18 |
|  | GUG | 1.21 | 1.21 | 1.21 | 1.21 | 1.24 | 1.21 | 1.21 | 1.21 | 1.21 | 1.18 | 1.24 | 1.24 | 1.18 |
| Serine(S) | UCU | 2.73 | 2.59 | 2.73 | 2.73 | 2.67 | 2.73 | 2.73 | 2.73 | 2.73 | 2.73 | 2.61 | 2.67 | 2.73 |
|  | UCC | 0.68 | 0.82 | 0.68 | 0.68 | 0.67 | 0.68 | 0.68 | 0.68 | 0.68 | 0.68 | 0.65 | 0.67 | 0.68 |
|  | UCA | 0.27 | 0.27 | 0.27 | 0.27 | 0.27 | 0.27 | 0.27 | 0.27 | 0.27 | 0.27 | 0.26 | 0.27 | 0.27 |
|  | UCG | 0.14 | 0.14 | 0.14 | 0.14 | 0.13 | 0.14 | 0.14 | 0.14 | 0.14 | 0.14 | 0.13 | 0.13 | 0.14 |
|  | AGU | 1.5 | 1.5 | 1.5 | 1.5 | 1.6 | 1.5 | 1.5 | 1.5 | 1.5 | 1.36 | 1.57 | 1.47 | 1.5 |
|  | AGC | 0.68 | 0.68 | 0.68 | 0.68 | 0.67 | 0.68 | 0.68 | 0.68 | 0.68 | 0.82 | 0.78 | 0.8 | 0.68 |
| Proline(P) | CCU | 2.09 | 2.09 | 2.04 | 2.09 | 2.14 | 2.09 | 2.09 | 2.04 | 2.09 | 2.04 | 2.09 | 2.09 | 2 |
|  | CCC | 0.73 | 0.73 | 0.8 | 0.73 | 0.65 | 0.73 | 0.73 | 0.8 | 0.73 | 0.71 | 0.73 | 0.73 | 0.73 |
|  | CCA | 1 | 1 | 0.98 | 1 | 1.12 | 1 | 1 | 0.98 | 1 | 1.07 | 1.09 | 1.09 | 1 |
|  | CCG | 0.18 | 0.18 | 0.18 | 0.18 | 0.09 | 0.18 | 0.18 | 0.18 | 0.18 | 0.18 | 0.09 | 0.09 | 0.27 |
| Threonine(T) | ACU | 1.78 | 1.78 | 1.67 | 1.78 | 1.84 | 1.67 | 1.84 | 1.67 | 1.89 | 1.78 | 1.89 | 1.89 | 1.78 |
|  | ACC | 0.89 | 0.89 | 0.89 | 0.89 | 0.86 | 0.89 | 0.86 | 0.89 | 0.89 | 0.89 | 0.78 | 0.78 | 0.89 |
|  | ACA | 1.11 | 1.11 | 1.11 | 1.11 | 1.08 | 1.11 | 1.08 | 1.11 | 1 | 1.11 | 1.11 | 1.11 | 1.11 |
|  | ACG | 0.22 | 0.22 | 0.33 | 0.22 | 0.22 | 0.33 | 0.22 | 0.33 | 0.22 | 0.22 | 0.22 | 0.22 | 0.22 |
| Alanine(A) | GCU | 1.94 | 1.94 | 1.94 | 1.94 | 2.06 | 1.94 | 1.94 | 1.94 | 1.94 | 1.87 | 2.06 | 2.06 | 1.87 |
|  | GCC | 1.03 | 1.03 | 1.03 | 1.03 | 0.9 | 1.03 | 1.03 | 1.03 | 1.03 | 1.07 | 0.9 | 0.9 | 1.07 |
|  | GCA | 1.03 | 1.03 | 1.03 | 1.03 | 1.03 | 1.03 | 1.03 | 1.03 | 1.03 | 1.07 | 1.03 | 1.03 | 1.07 |
|  | GCG | 0 | 0 | 0 | 0 | 0 | 0 | 0 | 0 | 0 | 0 | 0 | 0 | 0 |
| Tyrosine(Y) | UAU | 1.79 | 1.79 | 1.79 | 1.79 | 1.72 | 1.79 | 1.79 | 1.79 | 1.79 | 1.79 | 1.72 | 1.72 | 1.79 |
|  | UAC | 0.21 | 0.21 | 0.21 | 0.21 | 0.28 | 0.21 | 0.21 | 0.21 | 0.21 | 0.21 | 0.28 | 0.28 | 0.21 |
| Histidine(H) | CAU | 1.5 | 1.5 | 1.6 | 1.5 | 1.5 | 1.5 | 1.5 | 1.6 | 1.5 | 1.5 | 1.5 | 1.5 | 1.5 |
|  | CAC | 0.5 | 0.5 | 0.4 | 0.5 | 0.5 | 0.5 | 0.5 | 0.4 | 0.5 | 0.5 | 0.5 | 0.5 | 0.5 |
| Glutamine(Q) | CAA | 0.74 | 0.74 | 0.74 | 0.74 | 0.74 | 0.74 | 0.74 | 0.74 | 0.74 | 0.69 | 0.74 | 0.74 | 0.74 |
|  | CAG | 1.26 | 1.26 | 1.26 | 1.26 | 1.26 | 1.26 | 1.26 | 1.26 | 1.26 | 1.31 | 1.26 | 1.26 | 1.26 |
| Asparagine(N) | AAU | 1.57 | 1.57 | 1.57 | 1.57 | 1.64 | 1.57 | 1.55 | 1.57 | 1.57 | 1.57 | 1.64 | 1.64 | 1.57 |
|  | AAC | 0.43 | 0.43 | 0.43 | 0.43 | 0.36 | 0.43 | 0.45 | 0.43 | 0.43 | 0.43 | 0.36 | 0.36 | 0.43 |
| Lysine(K) | AAA | 0.96 | 1 | 0.96 | 0.96 | 1 | 0.96 | 0.96 | 0.89 | 0.96 | 0.96 | 1 | 0.93 | 0.96 |
|  | AAG | 1.04 | 1 | 1.04 | 1.04 | 1 | 1.04 | 1.04 | 1.11 | 1.04 | 1.04 | 1 | 1.07 | 1.04 |
| Aspartic acid(D) | GAU | 1.49 | 1.49 | 1.49 | 1.49 | 1.49 | 1.49 | 1.49 | 1.49 | 1.49 | 1.49 | 1.49 | 1.49 | 1.49 |
|  | GAC | 0.51 | 0.51 | 0.51 | 0.51 | 0.51 | 0.51 | 0.51 | 0.51 | 0.51 | 0.51 | 0.51 | 0.51 | 0.51 |
| Glutamic acid(E) | GAA | 1.29 | 1.29 | 1.29 | 1.29 | 1.43 | 1.29 | 1.29 | 1.29 | 1.29 | 1.29 | 1.43 | 1.43 | 1.29 |
|  | GAG | 0.71 | 0.71 | 0.71 | 0.71 | 0.57 | 0.71 | 0.71 | 0.71 | 0.71 | 0.71 | 0.57 | 0.57 | 0.71 |
| Cysteine C | UGU | 1.5 | 1.5 | 1.5 | 1.5 | 1.5 | 1.5 | 1.5 | 1.5 | 1.5 | 1.5 | 1.5 | 1.5 | 1.5 |
|  | UGC | 0.5 | 0.5 | 0.5 | 0.5 | 0.5 | 0.5 | 0.5 | 0.5 | 0.5 | 0.5 | 0.5 | 0.5 | 0.5 |
| Arginine R | CGU | 2.4 | 2.32 | 2.4 | 2.4 | 2.9 | 2.4 | 2.4 | 2.4 | 2.4 | 2.4 | 2.9 | 2.9 | 2.4 |
|  | CGC | 0.8 | 0.77 | 0.8 | 0.8 | 0.41 | 0.8 | 0.8 | 0.8 | 0.8 | 0.8 | 0.41 | 0.41 | 0.8 |
|  | CGA | 0 | 0 | 0 | 0 | 0 | 0 | 0 | 0 | 0 | 0 | 0 | 0 | 0 |
|  | CGG | 0.6 | 0.58 | 0.6 | 0.6 | 0.83 | 0.6 | 0.6 | 0.6 | 0.6 | 0.6 | 0.83 | 0.83 | 0.6 |
|  | AGA | 1 | 0.97 | 0.8 | 1 | 1.03 | 1 | 1.2 | 0.8 | 1 | 1 | 1.03 | 1.03 | 1.2 |
|  | AGG | 1.2 | 1.35 | 1.4 | 1.2 | 0.83 | 1.2 | 1 | 1.4 | 1.2 | 1.2 | 0.83 | 0.83 | 1 |
| Glycine (G) | GGU | 1.6 | 1.6 | 1.6 | 1.6 | 1.71 | 1.6 | 1.6 | 1.6 | 1.6 | 1.6 | 1.53 | 1.6 | 1.49 |
|  | GGC | 1.49 | 1.49 | 1.49 | 1.49 | 1.49 | 1.49 | 1.49 | 1.49 | 1.49 | 1.49 | 1.53 | 1.49 | 1.49 |
|  | GGA | 0.46 | 0.46 | 0.46 | 0.46 | 0.34 | 0.46 | 0.46 | 0.46 | 0.46 | 0.46 | 0.47 | 0.46 | 0.46 |
|  | GGG | 0.46 | 0.46 | 0.46 | 0.46 | 0.46 | 0.46 | 0.46 | 0.46 | 0.46 | 0.46 | 0.47 | 0.46 | 0.57 |

**Supplementary table 2.2:** Values of Relative Synonymous Codon Usage (RSCU) for accession number from KU707731.1 to KU707743.1 of L1 protein of Human Papillomavirus 18.

| **Amino acids** | **Codons** | **KU707731.1** | **KU707732.1** | **KU707733.1** | **KU707734.1** | **KU707735.1** | **KU707736.1** | **KU707737.1** | **KU707738.1** | **KU707739.1** | **KU707740.1** | **KU707741.1** | **KU707742.1** | **KU707743.1** |
| --- | --- | --- | --- | --- | --- | --- | --- | --- | --- | --- | --- | --- | --- | --- |
| Phenylalanine(F) | UUU | 1.75 | 1.83 | 1.75 | 1.83 | 1.83 | 1.83 | 1.83 | 1.83 | 1.83 | 1.83 | 1.83 | 1.83 | 1.83 |
|  | UUC | 0.25 | 0.17 | 0.25 | 0.17 | 0.17 | 0.17 | 0.17 | 0.17 | 0.17 | 0.17 | 0.17 | 0.17 | 0.17 |
| Leucine(L) | UUA | 2.82 | 2.88 | 2.82 | 2.94 | 2.94 | 2.94 | 2.94 | 2.82 | 2.71 | 2.88 | 2.88 | 2.88 | 2.82 |
|  | UUG | 1.59 | 1.56 | 1.59 | 1.53 | 1.53 | 1.53 | 1.53 | 1.65 | 1.65 | 1.56 | 1.56 | 1.56 | 1.65 |
|  | CUU | 0.61 | 0.6 | 0.61 | 0.59 | 0.59 | 0.59 | 0.59 | 0.59 | 0.59 | 0.6 | 0.6 | 0.6 | 0.59 |
|  | CUC | 0 | 0 | 0 | 0 | 0 | 0 | 0 | 0 | 0 | 0 | 0 | 0 | 0 |
|  | CUA | 0.49 | 0.48 | 0.49 | 0.47 | 0.47 | 0.47 | 0.47 | 0.47 | 0.59 | 0.48 | 0.48 | 0.48 | 0.47 |
|  | CUG | 0.49 | 0.48 | 0.49 | 0.47 | 0.47 | 0.47 | 0.47 | 0.47 | 0.47 | 0.48 | 0.48 | 0.48 | 0.47 |
| Isoleucine(I) | AUU | 2.4 | 2.4 | 2.4 | 2.31 | 2.31 | 2.31 | 2.31 | 2.4 | 2.31 | 2.4 | 2.4 | 2.4 | 2.4 |
|  | AUC | 0 | 0 | 0 | 0.12 | 0.12 | 0.12 | 0.12 | 0 | 0.12 | 0 | 0 | 0 | 0 |
|  | AUA | 0.6 | 0.6 | 0.6 | 0.58 | 0.58 | 0.58 | 0.58 | 0.6 | 0.58 | 0.6 | 0.6 | 0.6 | 0.6 |
| Valine(V) | GUU | 1.49 | 1.49 | 1.49 | 1.43 | 1.43 | 1.43 | 1.43 | 1.55 | 1.43 | 1.49 | 1.49 | 1.49 | 1.55 |
|  | GUC | 0.09 | 0.09 | 0.09 | 0.1 | 0.1 | 0.1 | 0.1 | 0.09 | 0.1 | 0.09 | 0.09 | 0.09 | 0.09 |
|  | GUA | 1.21 | 1.21 | 1.21 | 1.24 | 1.24 | 1.24 | 1.24 | 1.18 | 1.24 | 1.21 | 1.21 | 1.21 | 1.18 |
|  | GUG | 1.21 | 1.21 | 1.21 | 1.24 | 1.24 | 1.24 | 1.24 | 1.18 | 1.24 | 1.21 | 1.21 | 1.21 | 1.18 |
| Serine(S) | UCU | 2.73 | 2.73 | 2.73 | 2.67 | 2.67 | 2.67 | 2.67 | 2.73 | 2.67 | 2.73 | 2.73 | 2.73 | 2.73 |
|  | UCC | 0.68 | 0.68 | 0.68 | 0.67 | 0.67 | 0.67 | 0.67 | 0.68 | 0.67 | 0.68 | 0.68 | 0.68 | 0.68 |
|  | UCA | 0.27 | 0.27 | 0.27 | 0.27 | 0.27 | 0.27 | 0.27 | 0.27 | 0.27 | 0.27 | 0.27 | 0.27 | 0.27 |
|  | UCG | 0.14 | 0.14 | 0.14 | 0.13 | 0.13 | 0.13 | 0.13 | 0.14 | 0.13 | 0.14 | 0.14 | 0.14 | 0.14 |
|  | AGU | 1.5 | 1.5 | 1.5 | 1.6 | 1.6 | 1.6 | 1.6 | 1.5 | 1.6 | 1.5 | 1.5 | 1.5 | 1.5 |
|  | AGC | 0.68 | 0.68 | 0.68 | 0.67 | 0.67 | 0.67 | 0.67 | 0.68 | 0.67 | 0.68 | 0.68 | 0.68 | 0.68 |
| Proline(P) | CCU | 2.09 | 2.09 | 2.09 | 2.14 | 2.14 | 2.14 | 2.14 | 2.04 | 2.09 | 2.09 | 2.09 | 2.09 | 2.04 |
|  | CCC | 0.73 | 0.73 | 0.73 | 0.65 | 0.65 | 0.65 | 0.65 | 0.71 | 0.73 | 0.73 | 0.73 | 0.73 | 0.71 |
|  | CCA | 1 | 1 | 1 | 1.12 | 1.12 | 1.12 | 1.12 | 1.07 | 1.09 | 1 | 1 | 1 | 1.07 |
|  | CCG | 0.18 | 0.18 | 0.18 | 0.09 | 0.09 | 0.09 | 0.09 | 0.18 | 0.09 | 0.18 | 0.18 | 0.18 | 0.18 |
| Threonine(T) | ACU | 1.78 | 1.78 | 1.78 | 1.84 | 1.84 | 1.84 | 1.84 | 1.67 | 1.89 | 1.78 | 1.78 | 1.78 | 1.78 |
|  | ACC | 0.89 | 0.89 | 0.89 | 0.86 | 0.86 | 0.86 | 0.86 | 1 | 0.78 | 0.89 | 0.89 | 0.89 | 0.89 |
|  | ACA | 1.11 | 1.11 | 1.11 | 1.08 | 1.08 | 1.08 | 1.08 | 1.11 | 1.11 | 1.11 | 1.11 | 1.11 | 1.11 |
|  | ACG | 0.22 | 0.22 | 0.22 | 0.22 | 0.22 | 0.22 | 0.22 | 0.22 | 0.22 | 0.22 | 0.22 | 0.22 | 0.22 |
| Alanine(A) | GCU | 1.94 | 1.94 | 1.94 | 2.06 | 2.06 | 2.06 | 2.06 | 1.87 | 2.06 | 1.94 | 1.94 | 1.94 | 1.87 |
|  | GCC | 1.03 | 1.03 | 1.03 | 0.9 | 0.9 | 0.9 | 0.9 | 1.07 | 0.9 | 1.03 | 1.03 | 1.03 | 1.07 |
|  | GCA | 1.03 | 1.03 | 1.03 | 1.03 | 1.03 | 1.03 | 1.03 | 1.07 | 1.03 | 1.03 | 1.03 | 1.03 | 1.07 |
|  | GCG | 0 | 0 | 0 | 0 | 0 | 0 | 0 | 0 | 0 | 0 | 0 | 0 | 0 |
| Tyrosine(Y) | UAU | 1.79 | 1.79 | 1.79 | 1.72 | 1.72 | 1.67 | 1.72 | 1.79 | 1.72 | 1.79 | 1.79 | 1.79 | 1.79 |
|  | UAC | 0.21 | 0.21 | 0.21 | 0.28 | 0.28 | 0.33 | 0.28 | 0.21 | 0.28 | 0.21 | 0.21 | 0.21 | 0.21 |
| Histidine(H) | CAU | 1.5 | 1.5 | 1.5 | 1.5 | 1.5 | 1.6 | 1.5 | 1.5 | 1.5 | 1.5 | 1.5 | 1.5 | 1.5 |
|  | CAC | 0.5 | 0.5 | 0.5 | 0.5 | 0.5 | 0.4 | 0.5 | 0.5 | 0.5 | 0.5 | 0.5 | 0.5 | 0.5 |
| Glutamine(Q) | CAA | 0.74 | 0.81 | 0.74 | 0.74 | 0.74 | 0.74 | 0.74 | 0.69 | 0.74 | 0.74 | 0.74 | 0.74 | 0.69 |
|  | CAG | 1.26 | 1.19 | 1.26 | 1.26 | 1.26 | 1.26 | 1.26 | 1.31 | 1.26 | 1.26 | 1.26 | 1.26 | 1.31 |
| Asparagine(N) | AAU | 1.57 | 1.57 | 1.57 | 1.64 | 1.64 | 1.64 | 1.64 | 1.57 | 1.64 | 1.57 | 1.57 | 1.57 | 1.57 |
|  | AAC | 0.43 | 0.43 | 0.43 | 0.36 | 0.36 | 0.36 | 0.36 | 0.43 | 0.36 | 0.43 | 0.43 | 0.43 | 0.43 |
| Lysine(K) | AAA | 0.96 | 0.96 | 0.96 | 1 | 1 | 1.03 | 1 | 0.96 | 1 | 0.96 | 0.96 | 0.96 | 0.96 |
|  | AAG | 1.04 | 1.04 | 1.04 | 1 | 1 | 0.97 | 1 | 1.04 | 1 | 1.04 | 1.04 | 1.04 | 1.04 |
| Aspartic acid(D) | GAU | 1.49 | 1.49 | 1.49 | 1.49 | 1.49 | 1.49 | 1.49 | 1.49 | 1.49 | 1.49 | 1.49 | 1.49 | 1.49 |
|  | GAC | 0.51 | 0.51 | 0.51 | 0.51 | 0.51 | 0.51 | 0.51 | 0.51 | 0.51 | 0.51 | 0.51 | 0.51 | 0.51 |
| Glutamic acid(E) | GAA | 1.29 | 1.29 | 1.29 | 1.43 | 1.43 | 1.38 | 1.43 | 1.29 | 1.43 | 1.29 | 1.29 | 1.29 | 1.29 |
|  | GAG | 0.71 | 0.71 | 0.71 | 0.57 | 0.57 | 0.62 | 0.57 | 0.71 | 0.57 | 0.71 | 0.71 | 0.71 | 0.71 |
| Cysteine C | UGU | 1.5 | 1.5 | 1.53 | 1.5 | 1.5 | 1.5 | 1.5 | 1.5 | 1.5 | 1.5 | 1.5 | 1.5 | 1.5 |
|  | UGC | 0.5 | 0.5 | 0.47 | 0.5 | 0.5 | 0.5 | 0.5 | 0.5 | 0.5 | 0.5 | 0.5 | 0.5 | 0.5 |
| Arginine R | CGU | 2.4 | 2.4 | 2.4 | 2.9 | 2.9 | 2.9 | 2.9 | 2.4 | 2.9 | 2.4 | 2.4 | 2.4 | 2.4 |
|  | CGC | 0.8 | 0.8 | 0.8 | 0.41 | 0.41 | 0.41 | 0.41 | 0.8 | 0.41 | 0.8 | 0.8 | 0.8 | 0.8 |
|  | CGA | 0 | 0 | 0 | 0 | 0 | 0 | 0 | 0 | 0 | 0 | 0 | 0.2 | 0 |
|  | CGG | 0.6 | 0.6 | 0.6 | 0.83 | 0.83 | 0.83 | 0.83 | 0.6 | 0.83 | 0.6 | 0.6 | 0.6 | 0.6 |
|  | AGA | 1 | 1 | 1 | 1.03 | 1.03 | 1.03 | 1.03 | 1 | 1.03 | 1 | 1 | 0.8 | 1 |
|  | AGG | 1.2 | 1.2 | 1.2 | 0.83 | 0.83 | 0.83 | 0.83 | 1.2 | 0.83 | 1.2 | 1.2 | 1.2 | 1.2 |
| Glycine (G) | GGU | 1.6 | 1.6 | 1.6 | 1.71 | 1.71 | 1.71 | 1.71 | 1.6 | 1.71 | 1.6 | 1.6 | 1.6 | 1.6 |
|  | GGC | 1.49 | 1.49 | 1.49 | 1.49 | 1.49 | 1.49 | 1.49 | 1.49 | 1.49 | 1.49 | 1.49 | 1.49 | 1.49 |
|  | GGA | 0.46 | 0.46 | 0.46 | 0.34 | 0.34 | 0.34 | 0.34 | 0.46 | 0.34 | 0.46 | 0.46 | 0.46 | 0.46 |
|  | GGG | 0.46 | 0.46 | 0.46 | 0.46 | 0.46 | 0.46 | 0.46 | 0.46 | 0.46 | 0.46 | 0.46 | 0.46 | 0.46 |

**Supplementary table 2.3:** Values of Relative Synonymous Codon Usage (RSCU) for accession number from KU707744.1 to KU707756.1 of L1 protein of Human Papillomavirus 18.

| **Amino acids** | **Codons** | **KU707744.1** | **KU707745.1** | **KU707746.1** | **KU707747.1** | **KU707748.1** | **KU707749.1** | **KU707750.1** | **KU707751.1** | **KU707752.1** | **KU707753.1** | **KU707754.1** | **KU707755.1** | **KU707756.1** |
| --- | --- | --- | --- | --- | --- | --- | --- | --- | --- | --- | --- | --- | --- | --- |
| Phenylalanine(F) | UUU | 1.83 | 1.83 | 1.83 | 1.83 | 1.83 | 1.83 | 1.83 | 1.83 | 1.83 | 1.83 | 1.83 | 1.83 | 1.83 |
|  | UUC | 0.17 | 0.17 | 0.17 | 0.17 | 0.17 | 0.17 | 0.17 | 0.17 | 0.17 | 0.17 | 0.17 | 0.17 | 0.17 |
| Leucine(L) | UUA | 2.82 | 2.82 | 2.82 | 2.82 | 2.88 | 2.88 | 2.88 | 2.88 | 2.82 | 2.82 | 2.82 | 2.88 | 2.88 |
|  | UUG | 1.65 | 1.65 | 1.65 | 1.65 | 1.56 | 1.56 | 1.56 | 1.56 | 1.65 | 1.59 | 1.59 | 1.56 | 1.56 |
|  | CUU | 0.59 | 0.59 | 0.59 | 0.59 | 0.6 | 0.6 | 0.6 | 0.6 | 0.59 | 0.61 | 0.61 | 0.6 | 0.6 |
|  | CUC | 0 | 0 | 0 | 0 | 0 | 0 | 0 | 0 | 0 | 0 | 0 | 0 | 0 |
|  | CUA | 0.47 | 0.47 | 0.47 | 0.47 | 0.48 | 0.48 | 0.48 | 0.48 | 0.47 | 0.49 | 0.49 | 0.48 | 0.48 |
|  | CUG | 0.47 | 0.47 | 0.47 | 0.47 | 0.48 | 0.48 | 0.48 | 0.48 | 0.47 | 0.49 | 0.49 | 0.48 | 0.48 |
| Isoleucine(I) | AUU | 2.4 | 2.4 | 2.4 | 2.4 | 2.4 | 2.4 | 2.4 | 2.4 | 2.4 | 2.4 | 2.4 | 2.4 | 2.4 |
|  | AUC | 0 | 0 | 0 | 0 | 0 | 0 | 0 | 0 | 0 | 0 | 0 | 0 | 0 |
|  | AUA | 0.6 | 0.6 | 0.6 | 0.6 | 0.6 | 0.6 | 0.6 | 0.6 | 0.6 | 0.6 | 0.6 | 0.6 | 0.6 |
| Valine(V) | GUU | 1.49 | 1.49 | 1.49 | 1.49 | 1.49 | 1.49 | 1.49 | 1.49 | 1.55 | 1.49 | 1.49 | 1.49 | 1.49 |
|  | GUC | 0.09 | 0.09 | 0.09 | 0.09 | 0.09 | 0.09 | 0.09 | 0.09 | 0.09 | 0.09 | 0.09 | 0.09 | 0.09 |
|  | GUA | 1.21 | 1.21 | 1.21 | 1.21 | 1.21 | 1.21 | 1.21 | 1.21 | 1.18 | 1.21 | 1.21 | 1.21 | 1.21 |
|  | GUG | 1.21 | 1.21 | 1.21 | 1.21 | 1.21 | 1.21 | 1.21 | 1.21 | 1.18 | 1.21 | 1.21 | 1.21 | 1.21 |
| Serine(S) | UCU | 2.73 | 2.73 | 2.73 | 2.73 | 2.73 | 2.73 | 2.73 | 2.73 | 2.73 | 2.67 | 2.67 | 2.73 | 2.73 |
|  | UCC | 0.68 | 0.68 | 0.68 | 0.68 | 0.68 | 0.68 | 0.68 | 0.68 | 0.68 | 0.67 | 0.67 | 0.68 | 0.68 |
|  | UCA | 0.27 | 0.27 | 0.27 | 0.27 | 0.27 | 0.27 | 0.27 | 0.27 | 0.27 | 0.4 | 0.4 | 0.27 | 0.27 |
|  | UCG | 0.14 | 0.14 | 0.14 | 0.14 | 0.14 | 0.14 | 0.14 | 0.14 | 0.14 | 0.13 | 0.13 | 0.14 | 0.14 |
|  | AGU | 1.5 | 1.5 | 1.5 | 1.5 | 1.5 | 1.5 | 1.5 | 1.5 | 1.5 | 1.47 | 1.47 | 1.5 | 1.5 |
|  | AGC | 0.68 | 0.68 | 0.68 | 0.68 | 0.68 | 0.68 | 0.68 | 0.68 | 0.68 | 0.67 | 0.67 | 0.68 | 0.68 |
| Proline(P) | CCU | 2.09 | 2 | 2.09 | 2.09 | 2.09 | 2.09 | 2.09 | 2.09 | 2.04 | 2.09 | 2.09 | 2.09 | 2.09 |
|  | CCC | 0.73 | 0.73 | 0.73 | 0.73 | 0.73 | 0.73 | 0.73 | 0.73 | 0.71 | 0.73 | 0.73 | 0.73 | 0.73 |
|  | CCA | 1 | 1 | 1 | 1 | 1 | 1 | 1 | 1 | 1.07 | 1 | 1 | 1 | 1 |
|  | CCG | 0.18 | 0.27 | 0.18 | 0.18 | 0.18 | 0.18 | 0.18 | 0.18 | 0.18 | 0.18 | 0.18 | 0.18 | 0.18 |
| Threonine(T) | ACU | 1.78 | 1.84 | 1.84 | 1.78 | 1.78 | 1.78 | 1.78 | 1.78 | 1.78 | 1.78 | 1.78 | 1.78 | 1.71 |
|  | ACC | 0.89 | 0.86 | 0.86 | 0.89 | 0.89 | 0.89 | 0.89 | 0.89 | 0.89 | 0.89 | 0.89 | 0.89 | 0.91 |
|  | ACA | 1.11 | 1.08 | 1.08 | 1.11 | 1.11 | 1.11 | 1.11 | 1.11 | 1.11 | 1.11 | 1.11 | 1.11 | 1.14 |
|  | ACG | 0.22 | 0.22 | 0.22 | 0.22 | 0.22 | 0.22 | 0.22 | 0.22 | 0.22 | 0.22 | 0.22 | 0.22 | 0.23 |
| Alanine(A) | GCU | 1.94 | 1.94 | 1.94 | 1.94 | 1.94 | 1.94 | 1.94 | 1.94 | 1.87 | 1.94 | 1.94 | 1.94 | 2 |
|  | GCC | 1.03 | 1.03 | 1.03 | 1.03 | 1.03 | 1.03 | 1.03 | 1.03 | 1.07 | 1.03 | 1.03 | 1.03 | 1 |
|  | GCA | 1.03 | 1.03 | 1.03 | 1.03 | 1.03 | 1.03 | 1.03 | 1.03 | 1.07 | 1.03 | 1.03 | 1.03 | 1 |
|  | GCG | 0 | 0 | 0 | 0 | 0 | 0 | 0 | 0 | 0 | 0 | 0 | 0 | 0 |
| Tyrosine(Y) | UAU | 1.79 | 1.79 | 1.79 | 1.79 | 1.79 | 1.79 | 1.79 | 1.79 | 1.79 | 1.79 | 1.79 | 1.79 | 1.79 |
|  | UAC | 0.21 | 0.21 | 0.21 | 0.21 | 0.21 | 0.21 | 0.21 | 0.21 | 0.21 | 0.21 | 0.21 | 0.21 | 0.21 |
| Histidine(H) | CAU | 1.5 | 1.5 | 1.6 | 1.5 | 1.5 | 1.5 | 1.5 | 1.5 | 1.5 | 1.5 | 1.5 | 1.5 | 1.5 |
|  | CAC | 0.5 | 0.5 | 0.4 | 0.5 | 0.5 | 0.5 | 0.5 | 0.5 | 0.5 | 0.5 | 0.5 | 0.5 | 0.5 |
| Glutamine(Q) | CAA | 0.74 | 0.74 | 0.74 | 0.74 | 0.74 | 0.74 | 0.74 | 0.74 | 0.69 | 0.74 | 0.74 | 0.74 | 0.74 |
|  | CAG | 1.26 | 1.26 | 1.26 | 1.26 | 1.26 | 1.26 | 1.26 | 1.26 | 1.31 | 1.26 | 1.26 | 1.26 | 1.26 |
| Asparagine(N) | AAU | 1.57 | 1.55 | 1.55 | 1.57 | 1.57 | 1.57 | 1.57 | 1.57 | 1.57 | 1.57 | 1.57 | 1.57 | 1.57 |
|  | AAC | 0.43 | 0.45 | 0.45 | 0.43 | 0.43 | 0.43 | 0.43 | 0.43 | 0.43 | 0.43 | 0.43 | 0.43 | 0.43 |
| Lysine(K) | AAA | 0.96 | 0.96 | 0.96 | 0.96 | 0.96 | 0.96 | 0.96 | 0.96 | 0.96 | 1 | 1 | 0.96 | 0.96 |
|  | AAG | 1.04 | 1.04 | 1.04 | 1.04 | 1.04 | 1.04 | 1.04 | 1.04 | 1.04 | 1 | 1 | 1.04 | 1.04 |
| Aspartic acid(D) | GAU | 1.49 | 1.49 | 1.49 | 1.49 | 1.49 | 1.49 | 1.49 | 1.49 | 1.49 | 1.49 | 1.49 | 1.49 | 1.49 |
|  | GAC | 0.51 | 0.51 | 0.51 | 0.51 | 0.51 | 0.51 | 0.51 | 0.51 | 0.51 | 0.51 | 0.51 | 0.51 | 0.51 |
| Glutamic acid(E) | GAA | 1.29 | 1.29 | 1.29 | 1.29 | 1.29 | 1.29 | 1.29 | 1.29 | 1.29 | 1.29 | 1.29 | 1.29 | 1.29 |
|  | GAG | 0.71 | 0.71 | 0.71 | 0.71 | 0.71 | 0.71 | 0.71 | 0.71 | 0.71 | 0.71 | 0.71 | 0.71 | 0.71 |
| Cysteine C | UGU | 1.5 | 1.5 | 1.5 | 1.5 | 1.5 | 1.5 | 1.5 | 1.5 | 1.5 | 1.5 | 1.5 | 1.5 | 1.5 |
|  | UGC | 0.5 | 0.5 | 0.5 | 0.5 | 0.5 | 0.5 | 0.5 | 0.5 | 0.5 | 0.5 | 0.5 | 0.5 | 0.5 |
| Arginine R | CGU | 2.4 | 2.4 | 2.32 | 2.4 | 2.4 | 2.4 | 2.4 | 2.4 | 2.4 | 2.48 | 2.48 | 2.4 | 2.4 |
|  | CGC | 0.8 | 0.8 | 0.97 | 0.8 | 0.8 | 0.8 | 0.8 | 0.8 | 0.8 | 0.83 | 0.83 | 0.8 | 0.8 |
|  | CGA | 0 | 0 | 0 | 0 | 0 | 0 | 0 | 0 | 0 | 0 | 0 | 0 | 0 |
|  | CGG | 0.6 | 0.6 | 0.58 | 0.6 | 0.6 | 0.6 | 0.6 | 0.6 | 0.6 | 0.62 | 0.62 | 0.6 | 0.6 |
|  | AGA | 1 | 1.2 | 1.16 | 1.2 | 1 | 1 | 1 | 1 | 1 | 0.83 | 0.83 | 1 | 1 |
|  | AGG | 1.2 | 1 | 0.97 | 1 | 1.2 | 1.2 | 1.2 | 1.2 | 1.2 | 1.24 | 1.24 | 1.2 | 1.2 |
| Glycine (G) | GGU | 1.6 | 1.6 | 1.6 | 1.6 | 1.6 | 1.6 | 1.6 | 1.6 | 1.6 | 1.6 | 1.6 | 1.6 | 1.6 |
|  | GGC | 1.49 | 1.49 | 1.49 | 1.49 | 1.49 | 1.49 | 1.49 | 1.49 | 1.49 | 1.49 | 1.49 | 1.49 | 1.49 |
|  | GGA | 0.46 | 0.46 | 0.46 | 0.46 | 0.46 | 0.46 | 0.46 | 0.46 | 0.46 | 0.46 | 0.46 | 0.46 | 0.46 |
|  | GGG | 0.46 | 0.46 | 0.46 | 0.46 | 0.46 | 0.46 | 0.46 | 0.46 | 0.46 | 0.46 | 0.46 | 0.46 | 0.46 |

**Supplementary table 2.4:** Values of Relative Synonymous Codon Usage (RSCU) for accession number from KU707757.1 to KU707769.1 of L1 protein of Human Papillomavirus 18.

| **Amino acids** | **Codons** | **KU707757.1** | **KU707758.1** | **KU707759.1** | **KU707760.1** | **KU707761.1** | **KU707762.1** | **KU707763.1** | **KU707764.1** | **KU707765.1** | **KU707766.1** | **KU707767.1** | **KU707768.1** | **KU707769.1** |
| --- | --- | --- | --- | --- | --- | --- | --- | --- | --- | --- | --- | --- | --- | --- |
| Phenylalanine(F) | UUU | 1.83 | 1.83 | 1.83 | 1.83 | 1.83 | 1.83 | 1.83 | 1.83 | 1.83 | 1.75 | 1.83 | 1.83 | 1.83 |
|  | UUC | 0.17 | 0.17 | 0.17 | 0.17 | 0.17 | 0.17 | 0.17 | 0.17 | 0.17 | 0.25 | 0.17 | 0.17 | 0.17 |
| Leucine(L) | UUA | 2.82 | 2.82 | 2.82 | 2.88 | 2.94 | 2.94 | 2.88 | 2.94 | 2.88 | 2.82 | 2.88 | 2.88 | 2.82 |
|  | UUG | 1.59 | 1.65 | 1.65 | 1.56 | 1.53 | 1.53 | 1.56 | 1.53 | 1.56 | 1.59 | 1.56 | 1.56 | 1.65 |
|  | CUU | 0.61 | 0.59 | 0.59 | 0.6 | 0.59 | 0.59 | 0.6 | 0.59 | 0.6 | 0.61 | 0.6 | 0.6 | 0.59 |
|  | CUC | 0 | 0 | 0 | 0 | 0 | 0 | 0 | 0 | 0 | 0 | 0 | 0 | 0 |
|  | CUA | 0.49 | 0.47 | 0.47 | 0.48 | 0.47 | 0.47 | 0.48 | 0.47 | 0.48 | 0.49 | 0.48 | 0.48 | 0.47 |
|  | CUG | 0.49 | 0.47 | 0.47 | 0.48 | 0.47 | 0.47 | 0.48 | 0.47 | 0.48 | 0.49 | 0.48 | 0.48 | 0.47 |
| Isoleucine(I) | AUU | 2.4 | 2.4 | 2.4 | 2.4 | 2.31 | 2.4 | 2.4 | 2.31 | 2.4 | 2.4 | 2.4 | 2.4 | 2.19 |
|  | AUC | 0 | 0 | 0 | 0 | 0.12 | 0 | 0 | 0.12 | 0 | 0 | 0 | 0 | 0.12 |
|  | AUA | 0.6 | 0.6 | 0.6 | 0.6 | 0.58 | 0.6 | 0.6 | 0.58 | 0.6 | 0.6 | 0.6 | 0.6 | 0.69 |
| Valine(V) | GUU | 1.49 | 1.49 | 1.49 | 1.49 | 1.43 | 1.55 | 1.49 | 1.43 | 1.49 | 1.49 | 1.49 | 1.49 | 1.43 |
|  | GUC | 0.09 | 0.09 | 0.09 | 0.09 | 0.1 | 0.09 | 0.09 | 0.1 | 0.09 | 0.09 | 0.09 | 0.09 | 0.1 |
|  | GUA | 1.21 | 1.21 | 1.21 | 1.21 | 1.24 | 1.18 | 1.21 | 1.24 | 1.21 | 1.21 | 1.21 | 1.21 | 1.24 |
|  | GUG | 1.21 | 1.21 | 1.21 | 1.21 | 1.24 | 1.18 | 1.21 | 1.24 | 1.21 | 1.21 | 1.21 | 1.21 | 1.24 |
| Serine(S) | UCU | 2.67 | 2.73 | 2.73 | 2.73 | 2.67 | 2.73 | 2.73 | 2.67 | 2.73 | 2.73 | 2.73 | 2.73 | 2.67 |
|  | UCC | 0.67 | 0.68 | 0.68 | 0.68 | 0.67 | 0.68 | 0.68 | 0.67 | 0.68 | 0.68 | 0.68 | 0.68 | 0.67 |
|  | UCA | 0.4 | 0.27 | 0.27 | 0.27 | 0.27 | 0.27 | 0.27 | 0.27 | 0.27 | 0.27 | 0.27 | 0.27 | 0.27 |
|  | UCG | 0.13 | 0.14 | 0.14 | 0.14 | 0.13 | 0.14 | 0.14 | 0.13 | 0.14 | 0.14 | 0.14 | 0.14 | 0.13 |
|  | AGU | 1.47 | 1.5 | 1.5 | 1.5 | 1.6 | 1.5 | 1.5 | 1.6 | 1.5 | 1.5 | 1.5 | 1.5 | 1.47 |
|  | AGC | 0.67 | 0.68 | 0.68 | 0.68 | 0.67 | 0.68 | 0.68 | 0.67 | 0.68 | 0.68 | 0.68 | 0.68 | 0.8 |
| Proline(P) | CCU | 2.09 | 2.09 | 2.09 | 2.09 | 2.14 | 2.09 | 2.09 | 2.14 | 2.09 | 2.09 | 2.09 | 2.09 | 2.09 |
|  | CCC | 0.73 | 0.73 | 0.73 | 0.73 | 0.65 | 0.73 | 0.73 | 0.65 | 0.73 | 0.73 | 0.73 | 0.73 | 0.73 |
|  | CCA | 1 | 1 | 1 | 1 | 1.12 | 1 | 1 | 1.12 | 1 | 1 | 1 | 1 | 1.09 |
|  | CCG | 0.18 | 0.18 | 0.18 | 0.18 | 0.09 | 0.18 | 0.18 | 0.09 | 0.18 | 0.18 | 0.18 | 0.18 | 0.09 |
| Threonine(T) | ACU | 1.78 | 1.84 | 1.84 | 1.67 | 1.84 | 1.78 | 1.78 | 1.84 | 1.78 | 1.78 | 1.78 | 1.78 | 1.89 |
|  | ACC | 0.89 | 0.86 | 0.86 | 0.89 | 0.86 | 0.89 | 0.89 | 0.86 | 1 | 0.89 | 0.89 | 0.89 | 0.78 |
|  | ACA | 1.11 | 1.08 | 1.08 | 1.11 | 1.08 | 1.11 | 1.11 | 1.08 | 1 | 1.11 | 1.11 | 1.11 | 1.11 |
|  | ACG | 0.22 | 0.22 | 0.22 | 0.33 | 0.22 | 0.22 | 0.22 | 0.22 | 0.22 | 0.22 | 0.22 | 0.22 | 0.22 |
| Alanine(A) | GCU | 1.94 | 1.94 | 1.94 | 1.94 | 2.06 | 1.87 | 1.94 | 2.06 | 1.94 | 1.94 | 1.94 | 1.94 | 2.06 |
|  | GCC | 1.03 | 1.03 | 1.03 | 1.03 | 0.9 | 1.07 | 1.03 | 0.9 | 1.03 | 1.03 | 1.03 | 1.03 | 0.9 |
|  | GCA | 1.03 | 1.03 | 1.03 | 1.03 | 1.03 | 1.07 | 1.03 | 1.03 | 1.03 | 1.03 | 1.03 | 1.03 | 1.03 |
|  | GCG | 0 | 0 | 0 | 0 | 0 | 0 | 0 | 0 | 0 | 0 | 0 | 0 | 0 |
| Tyrosine(Y) | UAU | 1.79 | 1.79 | 1.79 | 1.79 | 1.72 | 1.79 | 1.79 | 1.72 | 1.79 | 1.79 | 1.79 | 1.79 | 1.72 |
|  | UAC | 0.21 | 0.21 | 0.21 | 0.21 | 0.28 | 0.21 | 0.21 | 0.28 | 0.21 | 0.21 | 0.21 | 0.21 | 0.28 |
| Histidine(H) | CAU | 1.5 | 1.5 | 1.5 | 1.5 | 1.5 | 1.5 | 1.5 | 1.5 | 1.5 | 1.5 | 1.5 | 1.5 | 1.5 |
|  | CAC | 0.5 | 0.5 | 0.5 | 0.5 | 0.5 | 0.5 | 0.5 | 0.5 | 0.5 | 0.5 | 0.5 | 0.5 | 0.5 |
| Glutamine(Q) | CAA | 0.74 | 0.74 | 0.74 | 0.74 | 0.74 | 0.74 | 0.74 | 0.74 | 0.74 | 0.74 | 0.74 | 0.74 | 0.74 |
|  | CAG | 1.26 | 1.26 | 1.26 | 1.26 | 1.26 | 1.26 | 1.26 | 1.26 | 1.26 | 1.26 | 1.26 | 1.26 | 1.26 |
| Asparagine(N) | AAU | 1.57 | 1.55 | 1.55 | 1.57 | 1.64 | 1.57 | 1.57 | 1.64 | 1.57 | 1.57 | 1.57 | 1.57 | 1.64 |
|  | AAC | 0.43 | 0.45 | 0.45 | 0.43 | 0.36 | 0.43 | 0.43 | 0.36 | 0.43 | 0.43 | 0.43 | 0.43 | 0.36 |
| Lysine(K) | AAA | 1 | 0.96 | 0.96 | 0.96 | 1 | 0.96 | 0.96 | 0.96 | 0.96 | 0.96 | 0.96 | 0.96 | 1 |
|  | AAG | 1 | 1.04 | 1.04 | 1.04 | 1 | 1.04 | 1.04 | 1.04 | 1.04 | 1.04 | 1.04 | 1.04 | 1 |
| Aspartic acid(D) | GAU | 1.49 | 1.49 | 1.49 | 1.49 | 1.49 | 1.49 | 1.49 | 1.49 | 1.49 | 1.49 | 1.49 | 1.49 | 1.49 |
|  | GAC | 0.51 | 0.51 | 0.51 | 0.51 | 0.51 | 0.51 | 0.51 | 0.51 | 0.51 | 0.51 | 0.51 | 0.51 | 0.51 |
| Glutamic acid(E) | GAA | 1.29 | 1.29 | 1.29 | 1.29 | 1.43 | 1.29 | 1.29 | 1.43 | 1.29 | 1.29 | 1.29 | 1.29 | 1.29 |
|  | GAG | 0.71 | 0.71 | 0.71 | 0.71 | 0.57 | 0.71 | 0.71 | 0.57 | 0.71 | 0.71 | 0.71 | 0.71 | 0.71 |
| Cysteine C | UGU | 1.5 | 1.5 | 1.5 | 1.5 | 1.5 | 1.5 | 1.5 | 1.5 | 1.5 | 1.5 | 1.5 | 1.5 | 1.5 |
|  | UGC | 0.5 | 0.5 | 0.5 | 0.5 | 0.5 | 0.5 | 0.5 | 0.5 | 0.5 | 0.5 | 0.5 | 0.5 | 0.5 |
| Arginine R | CGU | 2.48 | 2.4 | 2.4 | 2.4 | 2.9 | 2.4 | 2.4 | 2.8 | 2.4 | 2.4 | 2.4 | 2.4 | 2.9 |
|  | CGC | 0.83 | 0.8 | 0.8 | 0.8 | 0.41 | 0.8 | 0.8 | 0.4 | 0.8 | 0.8 | 0.8 | 0.8 | 0.41 |
|  | CGA | 0 | 0 | 0 | 0 | 0 | 0 | 0 | 0 | 0 | 0 | 0 | 0 | 0 |
|  | CGG | 0.62 | 0.6 | 0.6 | 0.6 | 0.83 | 0.6 | 0.6 | 0.8 | 0.6 | 0.6 | 0.6 | 0.6 | 0.83 |
|  | AGA | 0.83 | 1.2 | 1.2 | 0.8 | 1.03 | 1.2 | 1 | 1.2 | 1 | 1 | 1 | 1 | 1.03 |
|  | AGG | 1.24 | 1 | 1 | 1.4 | 0.83 | 1 | 1.2 | 0.8 | 1.2 | 1.2 | 1.2 | 1.2 | 0.83 |
| Glycine (G) | GGU | 1.6 | 1.6 | 1.6 | 1.6 | 1.71 | 1.49 | 1.6 | 1.71 | 1.6 | 1.6 | 1.6 | 1.6 | 1.6 |
|  | GGC | 1.49 | 1.49 | 1.49 | 1.49 | 1.49 | 1.49 | 1.49 | 1.49 | 1.49 | 1.49 | 1.49 | 1.49 | 1.49 |
|  | GGA | 0.46 | 0.46 | 0.46 | 0.46 | 0.34 | 0.46 | 0.46 | 0.34 | 0.46 | 0.46 | 0.46 | 0.46 | 0.46 |
|  | GGG | 0.46 | 0.46 | 0.46 | 0.46 | 0.46 | 0.57 | 0.46 | 0.46 | 0.46 | 0.46 | 0.46 | 0.46 | 0.46 |

**Supplementary table 2.5:** Values of Relative Synonymous Codon Usage (RSCU) for accession number from KU707770.1 to KU707782.1 of L1 protein of Human Papillomavirus 18.

| **Amino acids** | **Codons** | **KU707770.1** | **KU707771.1** | **KU707772.1** | **KU707773.1** | **KU707774.1** | **KU707775.1** | **KU707776.1** | **KU707777.1** | **KU707778.1** | **KU707779.1** | **KU707780.1** | **KU707781.1** | **KU707782.1** |
| --- | --- | --- | --- | --- | --- | --- | --- | --- | --- | --- | --- | --- | --- | --- |
| Phenylalanine(F) | UUU | 1.75 | 1.83 | 1.83 | 1.83 | 1.75 | 1.83 | 1.83 | 1.83 | 1.83 | 1.83 | 1.83 | 1.83 | 1.83 |
|  | UUC | 0.25 | 0.17 | 0.17 | 0.17 | 0.25 | 0.17 | 0.17 | 0.17 | 0.17 | 0.17 | 0.17 | 0.17 | 0.17 |
| Leucine(L) | UUA | 2.82 | 2.82 | 2.88 | 2.88 | 2.82 | 2.88 | 2.88 | 2.94 | 2.82 | 2.88 | 2.82 | 2.94 | 2.88 |
|  | UUG | 1.59 | 1.65 | 1.56 | 1.56 | 1.59 | 1.56 | 1.56 | 1.53 | 1.65 | 1.56 | 1.65 | 1.53 | 1.56 |
|  | CUU | 0.61 | 0.59 | 0.6 | 0.6 | 0.61 | 0.6 | 0.6 | 0.59 | 0.59 | 0.6 | 0.59 | 0.59 | 0.6 |
|  | CUC | 0 | 0 | 0 | 0 | 0 | 0 | 0 | 0 | 0 | 0 | 0 | 0 | 0 |
|  | CUA | 0.49 | 0.47 | 0.48 | 0.48 | 0.49 | 0.48 | 0.48 | 0.47 | 0.47 | 0.48 | 0.47 | 0.47 | 0.48 |
|  | CUG | 0.49 | 0.47 | 0.48 | 0.48 | 0.49 | 0.48 | 0.48 | 0.47 | 0.47 | 0.48 | 0.47 | 0.47 | 0.48 |
| Isoleucine(I) | AUU | 2.4 | 2.4 | 2.4 | 2.4 | 2.4 | 2.4 | 2.4 | 2.4 | 2.4 | 2.4 | 2.31 | 2.31 | 2.4 |
|  | AUC | 0 | 0 | 0 | 0 | 0 | 0 | 0 | 0 | 0 | 0 | 0.12 | 0.12 | 0 |
|  | AUA | 0.6 | 0.6 | 0.6 | 0.6 | 0.6 | 0.6 | 0.6 | 0.6 | 0.6 | 0.6 | 0.58 | 0.58 | 0.6 |
| Valine(V) | GUU | 1.49 | 1.49 | 1.49 | 1.49 | 1.49 | 1.49 | 1.49 | 1.55 | 1.49 | 1.49 | 1.43 | 1.43 | 1.49 |
|  | GUC | 0.09 | 0.09 | 0.09 | 0.09 | 0.09 | 0.09 | 0.09 | 0.09 | 0.09 | 0.09 | 0.1 | 0.1 | 0.09 |
|  | GUA | 1.21 | 1.21 | 1.21 | 1.21 | 1.21 | 1.21 | 1.21 | 1.18 | 1.21 | 1.21 | 1.24 | 1.24 | 1.21 |
|  | GUG | 1.21 | 1.21 | 1.21 | 1.21 | 1.21 | 1.21 | 1.21 | 1.18 | 1.21 | 1.21 | 1.24 | 1.24 | 1.21 |
| Serine(S) | UCU | 2.73 | 2.73 | 2.73 | 2.73 | 2.73 | 2.73 | 2.73 | 2.73 | 2.73 | 2.73 | 2.67 | 2.67 | 2.73 |
|  | UCC | 0.68 | 0.68 | 0.68 | 0.68 | 0.68 | 0.68 | 0.68 | 0.68 | 0.68 | 0.68 | 0.67 | 0.67 | 0.68 |
|  | UCA | 0.27 | 0.27 | 0.27 | 0.27 | 0.27 | 0.27 | 0.27 | 0.27 | 0.27 | 0.27 | 0.27 | 0.27 | 0.27 |
|  | UCG | 0.14 | 0.14 | 0.14 | 0.14 | 0.14 | 0.14 | 0.14 | 0.14 | 0.14 | 0.14 | 0.13 | 0.13 | 0.14 |
|  | AGU | 1.5 | 1.5 | 1.5 | 1.5 | 1.5 | 1.5 | 1.5 | 1.5 | 1.5 | 1.5 | 1.6 | 1.6 | 1.5 |
|  | AGC | 0.68 | 0.68 | 0.68 | 0.68 | 0.68 | 0.68 | 0.68 | 0.68 | 0.68 | 0.68 | 0.67 | 0.67 | 0.68 |
| Proline(P) | CCU | 2.09 | 2.13 | 2.09 | 2.09 | 2.09 | 2.09 | 2.09 | 2.09 | 2.09 | 2.04 | 2.09 | 2.09 | 2.09 |
|  | CCC | 0.73 | 0.71 | 0.73 | 0.73 | 0.73 | 0.73 | 0.73 | 0.73 | 0.73 | 0.8 | 0.73 | 0.64 | 0.73 |
|  | CCA | 1 | 0.98 | 1 | 1 | 1 | 1 | 1 | 1 | 1 | 0.98 | 1.09 | 1.18 | 1 |
|  | CCG | 0.18 | 0.18 | 0.18 | 0.18 | 0.18 | 0.18 | 0.18 | 0.18 | 0.18 | 0.18 | 0.09 | 0.09 | 0.18 |
| Threonine(T) | ACU | 1.78 | 1.84 | 1.78 | 1.78 | 1.78 | 1.78 | 1.78 | 1.78 | 1.84 | 1.78 | 1.89 | 1.84 | 1.78 |
|  | ACC | 0.89 | 0.86 | 0.89 | 0.89 | 0.89 | 0.89 | 0.89 | 0.89 | 0.86 | 0.89 | 0.78 | 0.86 | 0.89 |
|  | ACA | 1.11 | 1.08 | 1.11 | 1.11 | 1.11 | 1.11 | 1.11 | 1.11 | 1.08 | 1.11 | 1.11 | 1.08 | 1.11 |
|  | ACG | 0.22 | 0.22 | 0.22 | 0.22 | 0.22 | 0.22 | 0.22 | 0.22 | 0.22 | 0.22 | 0.22 | 0.22 | 0.22 |
| Alanine(A) | GCU | 1.94 | 1.94 | 1.94 | 1.94 | 1.94 | 1.94 | 1.94 | 1.87 | 1.94 | 1.94 | 2.06 | 2.06 | 1.94 |
|  | GCC | 1.03 | 1.03 | 1.03 | 1.03 | 1.03 | 1.03 | 1.03 | 1.07 | 1.03 | 1.03 | 0.9 | 0.9 | 1.03 |
|  | GCA | 1.03 | 1.03 | 1.03 | 1.03 | 1.03 | 1.03 | 1.03 | 1.07 | 1.03 | 1.03 | 1.03 | 1.03 | 1.03 |
|  | GCG | 0 | 0 | 0 | 0 | 0 | 0 | 0 | 0 | 0 | 0 | 0 | 0 | 0 |
| Tyrosine(Y) | UAU | 1.79 | 1.79 | 1.79 | 1.79 | 1.79 | 1.79 | 1.79 | 1.79 | 1.79 | 1.79 | 1.72 | 1.72 | 1.79 |
|  | UAC | 0.21 | 0.21 | 0.21 | 0.21 | 0.21 | 0.21 | 0.21 | 0.21 | 0.21 | 0.21 | 0.28 | 0.28 | 0.21 |
| Histidine(H) | CAU | 1.5 | 1.47 | 1.5 | 1.5 | 1.5 | 1.5 | 1.5 | 1.5 | 1.5 | 1.6 | 1.5 | 1.53 | 1.5 |
|  | CAC | 0.5 | 0.53 | 0.5 | 0.5 | 0.5 | 0.5 | 0.5 | 0.5 | 0.5 | 0.4 | 0.5 | 0.47 | 0.5 |
| Glutamine(Q) | CAA | 0.74 | 0.74 | 0.74 | 0.74 | 0.74 | 0.74 | 0.74 | 0.74 | 0.74 | 0.74 | 0.74 | 0.69 | 0.74 |
|  | CAG | 1.26 | 1.26 | 1.26 | 1.26 | 1.26 | 1.26 | 1.26 | 1.26 | 1.26 | 1.26 | 1.26 | 1.31 | 1.26 |
| Asparagine(N) | AAU | 1.57 | 1.55 | 1.57 | 1.57 | 1.57 | 1.57 | 1.57 | 1.57 | 1.55 | 1.57 | 1.64 | 1.62 | 1.57 |
|  | AAC | 0.43 | 0.45 | 0.43 | 0.43 | 0.43 | 0.43 | 0.43 | 0.43 | 0.45 | 0.43 | 0.36 | 0.38 | 0.43 |
| Lysine(K) | AAA | 0.96 | 0.96 | 0.96 | 0.96 | 0.96 | 0.96 | 0.96 | 0.96 | 0.96 | 0.96 | 1 | 1 | 0.96 |
|  | AAG | 1.04 | 1.04 | 1.04 | 1.04 | 1.04 | 1.04 | 1.04 | 1.04 | 1.04 | 1.04 | 1 | 1 | 1.04 |
| Aspartic acid(D) | GAU | 1.49 | 1.49 | 1.49 | 1.49 | 1.49 | 1.49 | 1.49 | 1.49 | 1.49 | 1.49 | 1.49 | 1.49 | 1.49 |
|  | GAC | 0.51 | 0.51 | 0.51 | 0.51 | 0.51 | 0.51 | 0.51 | 0.51 | 0.51 | 0.51 | 0.51 | 0.51 | 0.51 |
| Glutamic acid(E) | GAA | 1.29 | 1.29 | 1.29 | 1.29 | 1.29 | 1.29 | 1.29 | 1.29 | 1.29 | 1.29 | 1.43 | 1.43 | 1.29 |
|  | GAG | 0.71 | 0.71 | 0.71 | 0.71 | 0.71 | 0.71 | 0.71 | 0.71 | 0.71 | 0.71 | 0.57 | 0.57 | 0.71 |
| Cysteine C | UGU | 1.53 | 1.5 | 1.5 | 1.5 | 1.5 | 1.5 | 1.5 | 1.5 | 1.5 | 1.5 | 1.5 | 1.5 | 1.5 |
|  | UGC | 0.47 | 0.5 | 0.5 | 0.5 | 0.5 | 0.5 | 0.5 | 0.5 | 0.5 | 0.5 | 0.5 | 0.5 | 0.5 |
| Arginine R | CGU | 2.4 | 2.4 | 2.4 | 2.4 | 2.4 | 2.4 | 2.4 | 2.4 | 2.4 | 2.4 | 2.9 | 2.9 | 2.4 |
|  | CGC | 0.8 | 0.8 | 0.8 | 0.8 | 0.8 | 0.8 | 0.8 | 0.8 | 0.8 | 0.8 | 0.41 | 0.41 | 0.8 |
|  | CGA | 0 | 0 | 0 | 0 | 0 | 0 | 0 | 0 | 0 | 0 | 0 | 0 | 0 |
|  | CGG | 0.6 | 0.6 | 0.6 | 0.6 | 0.6 | 0.6 | 0.6 | 0.6 | 0.6 | 0.6 | 0.83 | 0.83 | 0.6 |
|  | AGA | 1 | 1.2 | 1 | 1 | 1 | 1 | 1.2 | 1.2 | 1.2 | 1 | 1.03 | 1.03 | 1 |
|  | AGG | 1.2 | 1 | 1.2 | 1.2 | 1.2 | 1.2 | 1 | 1 | 1 | 1.2 | 0.83 | 0.83 | 1.2 |
| Glycine (G) | GGU | 1.6 | 1.6 | 1.6 | 1.6 | 1.6 | 1.6 | 1.6 | 1.49 | 1.6 | 1.6 | 1.71 | 1.71 | 1.6 |
|  | GGC | 1.49 | 1.49 | 1.49 | 1.49 | 1.49 | 1.49 | 1.49 | 1.49 | 1.49 | 1.49 | 1.49 | 1.49 | 1.49 |
|  | GGA | 0.46 | 0.46 | 0.46 | 0.46 | 0.46 | 0.46 | 0.46 | 0.46 | 0.46 | 0.46 | 0.34 | 0.34 | 0.46 |
|  | GGG | 0.46 | 0.46 | 0.46 | 0.46 | 0.46 | 0.46 | 0.46 | 0.57 | 0.46 | 0.46 | 0.46 | 0.46 | 0.46 |

**Supplementary table 2.6:** Values of Relative Synonymous Codon Usage (RSCU) for accession number from KU707783.1 to KU707795.1 of L1 protein of Human Papillomavirus 18.

| **Amino acids** | **Codons** | **KU707783.1** | **KU707784.1** | **KU707785.1** | **KU707786.1** | **KU707787.1** | **KU707788.1** | **KU707789.1** | **KU707790.1** | **KU707791.1** | **KU707792.1** | **KU707793.1** | **KU707794.1** | **KU707795.1** |
| --- | --- | --- | --- | --- | --- | --- | --- | --- | --- | --- | --- | --- | --- | --- |
| Phenylalanine(F) | UUU | 1.83 | 1.83 | 1.83 | 1.83 | 1.83 | 1.83 | 1.83 | 1.83 | 1.83 | 1.83 | 1.83 | 1.83 | 1.75 |
|  | UUC | 0.17 | 0.17 | 0.17 | 0.17 | 0.17 | 0.17 | 0.17 | 0.17 | 0.17 | 0.17 | 0.17 | 0.17 | 0.25 |
| Leucine(L) | UUA | 2.88 | 2.82 | 2.88 | 2.82 | 2.88 | 2.82 | 2.82 | 2.88 | 2.82 | 2.88 | 2.88 | 2.88 | 2.82 |
|  | UUG | 1.56 | 1.65 | 1.56 | 1.65 | 1.56 | 1.65 | 1.65 | 1.56 | 1.65 | 1.56 | 1.56 | 1.56 | 1.59 |
|  | CUU | 0.6 | 0.59 | 0.6 | 0.59 | 0.6 | 0.59 | 0.59 | 0.6 | 0.59 | 0.6 | 0.6 | 0.6 | 0.61 |
|  | CUC | 0 | 0 | 0 | 0 | 0 | 0 | 0 | 0 | 0 | 0 | 0 | 0 | 0 |
|  | CUA | 0.48 | 0.47 | 0.48 | 0.47 | 0.48 | 0.47 | 0.47 | 0.48 | 0.47 | 0.48 | 0.48 | 0.48 | 0.49 |
|  | CUG | 0.48 | 0.47 | 0.48 | 0.47 | 0.48 | 0.47 | 0.47 | 0.48 | 0.47 | 0.48 | 0.48 | 0.48 | 0.49 |
| Isoleucine(I) | AUU | 2.4 | 2.4 | 2.4 | 2.4 | 2.4 | 2.4 | 2.4 | 2.4 | 2.4 | 2.4 | 2.4 | 2.4 | 2.4 |
|  | AUC | 0 | 0 | 0 | 0 | 0 | 0 | 0 | 0 | 0 | 0 | 0 | 0 | 0 |
|  | AUA | 0.6 | 0.6 | 0.6 | 0.6 | 0.6 | 0.6 | 0.6 | 0.6 | 0.6 | 0.6 | 0.6 | 0.6 | 0.6 |
| Valine(V) | GUU | 1.49 | 1.49 | 1.49 | 1.49 | 1.49 | 1.55 | 1.49 | 1.49 | 1.49 | 1.49 | 1.49 | 1.49 | 1.49 |
|  | GUC | 0.09 | 0.09 | 0.09 | 0.09 | 0.09 | 0.09 | 0.09 | 0.09 | 0.09 | 0.09 | 0.09 | 0.09 | 0.09 |
|  | GUA | 1.21 | 1.21 | 1.21 | 1.21 | 1.21 | 1.18 | 1.21 | 1.21 | 1.21 | 1.21 | 1.21 | 1.21 | 1.21 |
|  | GUG | 1.21 | 1.21 | 1.21 | 1.21 | 1.21 | 1.18 | 1.21 | 1.21 | 1.21 | 1.21 | 1.21 | 1.21 | 1.21 |
| Serine(S) | UCU | 2.73 | 2.73 | 2.73 | 2.65 | 2.73 | 2.73 | 2.73 | 2.73 | 2.73 | 2.73 | 2.73 | 2.73 | 2.73 |
|  | UCC | 0.68 | 0.68 | 0.68 | 0.7 | 0.68 | 0.68 | 0.68 | 0.68 | 0.68 | 0.68 | 0.68 | 0.68 | 0.68 |
|  | UCA | 0.27 | 0.27 | 0.27 | 0.28 | 0.27 | 0.27 | 0.27 | 0.27 | 0.27 | 0.27 | 0.27 | 0.27 | 0.27 |
|  | UCG | 0.14 | 0.14 | 0.14 | 0.28 | 0.14 | 0.14 | 0.14 | 0.14 | 0.14 | 0.14 | 0.14 | 0.14 | 0.14 |
|  | AGU | 1.5 | 1.5 | 1.5 | 1.4 | 1.5 | 1.5 | 1.5 | 1.5 | 1.5 | 1.5 | 1.5 | 1.5 | 1.5 |
|  | AGC | 0.68 | 0.68 | 0.68 | 0.7 | 0.68 | 0.68 | 0.68 | 0.68 | 0.68 | 0.68 | 0.68 | 0.68 | 0.68 |
| Proline(P) | CCU | 2.09 | 2.09 | 2.09 | 2.09 | 2.09 | 2.04 | 2.09 | 2.09 | 2.09 | 2.09 | 2.09 | 2.09 | 2.09 |
|  | CCC | 0.73 | 0.73 | 0.73 | 0.73 | 0.73 | 0.71 | 0.73 | 0.73 | 0.73 | 0.73 | 0.73 | 0.73 | 0.73 |
|  | CCA | 1 | 1 | 1 | 1 | 1 | 1.07 | 1 | 1 | 1 | 1 | 1 | 1 | 1 |
|  | CCG | 0.18 | 0.18 | 0.18 | 0.18 | 0.18 | 0.18 | 0.18 | 0.18 | 0.18 | 0.18 | 0.18 | 0.18 | 0.18 |
| Threonine(T) | ACU | 1.89 | 1.84 | 1.89 | 1.84 | 1.78 | 1.67 | 1.84 | 1.78 | 1.84 | 1.78 | 1.89 | 1.78 | 1.78 |
|  | ACC | 0.89 | 0.86 | 0.89 | 0.86 | 0.89 | 1 | 0.86 | 0.89 | 0.86 | 0.89 | 0.89 | 0.89 | 0.89 |
|  | ACA | 1 | 1.08 | 1 | 1.08 | 1.11 | 1.11 | 1.08 | 1.11 | 1.08 | 1.11 | 1 | 1.11 | 1.11 |
|  | ACG | 0.22 | 0.22 | 0.22 | 0.22 | 0.22 | 0.22 | 0.22 | 0.22 | 0.22 | 0.22 | 0.22 | 0.22 | 0.22 |
| Alanine(A) | GCU | 1.94 | 1.94 | 1.94 | 1.94 | 1.94 | 1.87 | 1.94 | 1.94 | 1.94 | 1.94 | 1.94 | 1.94 | 1.94 |
|  | GCC | 1.03 | 1.03 | 1.03 | 1.03 | 1.03 | 1.07 | 1.03 | 1.03 | 1.03 | 1.03 | 1.03 | 1.03 | 1.03 |
|  | GCA | 1.03 | 1.03 | 1.03 | 1.03 | 1.03 | 1.07 | 1.03 | 1.03 | 1.03 | 1.03 | 1.03 | 1.03 | 1.03 |
|  | GCG | 0 | 0 | 0 | 0 | 0 | 0 | 0 | 0 | 0 | 0 | 0 | 0 | 0 |
| Tyrosine(Y) | UAU | 1.79 | 1.79 | 1.79 | 1.79 | 1.79 | 1.79 | 1.79 | 1.79 | 1.79 | 1.79 | 1.79 | 1.79 | 1.79 |
|  | UAC | 0.21 | 0.21 | 0.21 | 0.21 | 0.21 | 0.21 | 0.21 | 0.21 | 0.21 | 0.21 | 0.21 | 0.21 | 0.21 |
| Histidine(H) | CAU | 1.5 | 1.5 | 1.5 | 1.5 | 1.5 | 1.5 | 1.5 | 1.5 | 1.5 | 1.5 | 1.5 | 1.5 | 1.5 |
|  | CAC | 0.5 | 0.5 | 0.5 | 0.5 | 0.5 | 0.5 | 0.5 | 0.5 | 0.5 | 0.5 | 0.5 | 0.5 | 0.5 |
| Glutamine(Q) | CAA | 0.74 | 0.74 | 0.74 | 0.74 | 0.74 | 0.69 | 0.74 | 0.74 | 0.74 | 0.74 | 0.74 | 0.74 | 0.74 |
|  | CAG | 1.26 | 1.26 | 1.26 | 1.26 | 1.26 | 1.31 | 1.26 | 1.26 | 1.26 | 1.26 | 1.26 | 1.26 | 1.26 |
| Asparagine(N) | AAU | 1.57 | 1.55 | 1.57 | 1.57 | 1.57 | 1.57 | 1.55 | 1.57 | 1.55 | 1.57 | 1.57 | 1.57 | 1.57 |
|  | AAC | 0.43 | 0.45 | 0.43 | 0.43 | 0.43 | 0.43 | 0.45 | 0.43 | 0.45 | 0.43 | 0.43 | 0.43 | 0.43 |
| Lysine(K) | AAA | 0.96 | 0.96 | 0.96 | 0.96 | 0.96 | 0.96 | 0.96 | 0.96 | 0.96 | 0.96 | 0.96 | 0.96 | 0.96 |
|  | AAG | 1.04 | 1.04 | 1.04 | 1.04 | 1.04 | 1.04 | 1.04 | 1.04 | 1.04 | 1.04 | 1.04 | 1.04 | 1.04 |
| Aspartic acid(D) | GAU | 1.49 | 1.49 | 1.49 | 1.49 | 1.49 | 1.49 | 1.49 | 1.49 | 1.49 | 1.49 | 1.49 | 1.49 | 1.49 |
|  | GAC | 0.51 | 0.51 | 0.51 | 0.51 | 0.51 | 0.51 | 0.51 | 0.51 | 0.51 | 0.51 | 0.51 | 0.51 | 0.51 |
| Glutamic acid(E) | GAA | 1.29 | 1.29 | 1.29 | 1.29 | 1.29 | 1.29 | 1.29 | 1.29 | 1.29 | 1.29 | 1.29 | 1.29 | 1.29 |
|  | GAG | 0.71 | 0.71 | 0.71 | 0.71 | 0.71 | 0.71 | 0.71 | 0.71 | 0.71 | 0.71 | 0.71 | 0.71 | 0.71 |
| Cysteine C | UGU | 1.5 | 1.5 | 1.5 | 1.5 | 1.5 | 1.5 | 1.5 | 1.5 | 1.5 | 1.5 | 1.5 | 1.5 | 1.5 |
|  | UGC | 0.5 | 0.5 | 0.5 | 0.5 | 0.5 | 0.5 | 0.5 | 0.5 | 0.5 | 0.5 | 0.5 | 0.5 | 0.5 |
| Arginine R | CGU | 2.4 | 2.4 | 2.4 | 2.4 | 2.4 | 2.4 | 2.4 | 2.4 | 2.4 | 2.4 | 2.4 | 2.4 | 2.4 |
|  | CGC | 0.8 | 0.8 | 0.8 | 0.8 | 0.8 | 0.8 | 0.8 | 0.8 | 0.8 | 0.8 | 0.8 | 0.8 | 0.8 |
|  | CGA | 0 | 0 | 0 | 0.2 | 0 | 0 | 0 | 0 | 0 | 0 | 0 | 0 | 0 |
|  | CGG | 0.6 | 0.6 | 0.6 | 0.6 | 0.6 | 0.6 | 0.6 | 0.6 | 0.6 | 0.6 | 0.6 | 0.6 | 0.6 |
|  | AGA | 1 | 1.2 | 1 | 1 | 1 | 1 | 1.2 | 1 | 1.2 | 1 | 1 | 1 | 1 |
|  | AGG | 1.2 | 1 | 1.2 | 1 | 1.2 | 1.2 | 1 | 1.2 | 1 | 1.2 | 1.2 | 1.2 | 1.2 |
| Glycine (G) | GGU | 1.6 | 1.6 | 1.6 | 1.6 | 1.6 | 1.6 | 1.6 | 1.6 | 1.6 | 1.6 | 1.6 | 1.6 | 1.6 |
|  | GGC | 1.49 | 1.49 | 1.49 | 1.49 | 1.49 | 1.49 | 1.49 | 1.49 | 1.49 | 1.49 | 1.49 | 1.49 | 1.49 |
|  | GGA | 0.46 | 0.46 | 0.46 | 0.46 | 0.46 | 0.46 | 0.46 | 0.46 | 0.46 | 0.46 | 0.46 | 0.46 | 0.46 |
|  | GGG | 0.46 | 0.46 | 0.46 | 0.46 | 0.46 | 0.46 | 0.46 | 0.46 | 0.46 | 0.46 | 0.46 | 0.46 | 0.46 |

**Supplementary table 2.7:** Values of Relative Synonymous Codon Usage (RSCU) for accession number from KU707796.1 to KU707808.1 of L1 protein of Human Papillomavirus 18.

| **Amino acids** | **Codons** | **KU707796.1** | **KU707797.1** | **KU707798.1** | **KU707799.1** | **KU707800.1** | **KU707801.1** | **KU707802.1** | **KU707803.1** | **KU707804.1** | **KU707805.1** | **KU707806.1** | **KU707807.1** | **KU707808.1** |
| --- | --- | --- | --- | --- | --- | --- | --- | --- | --- | --- | --- | --- | --- | --- |
| Phenylalanine(F) | UUU | 1.83 | 1.83 | 1.83 | 1.83 | 1.83 | 1.83 | 1.83 | 1.83 | 1.83 | 1.83 | 1.83 | 1.83 | 1.83 |
|  | UUC | 0.17 | 0.17 | 0.17 | 0.17 | 0.17 | 0.17 | 0.17 | 0.17 | 0.17 | 0.17 | 0.17 | 0.17 | 0.17 |
| Leucine(L) | UUA | 2.88 | 2.88 | 2.88 | 2.88 | 2.88 | 2.94 | 2.94 | 2.88 | 2.88 | 2.94 | 2.88 | 2.82 | 2.88 |
|  | UUG | 1.56 | 1.56 | 1.56 | 1.56 | 1.56 | 1.53 | 1.53 | 1.56 | 1.56 | 1.53 | 1.56 | 1.65 | 1.56 |
|  | CUU | 0.6 | 0.6 | 0.6 | 0.6 | 0.6 | 0.59 | 0.59 | 0.6 | 0.6 | 0.59 | 0.6 | 0.59 | 0.6 |
|  | CUC | 0 | 0 | 0 | 0 | 0 | 0 | 0 | 0 | 0 | 0 | 0 | 0 | 0 |
|  | CUA | 0.48 | 0.48 | 0.48 | 0.48 | 0.48 | 0.47 | 0.47 | 0.48 | 0.48 | 0.47 | 0.48 | 0.47 | 0.48 |
|  | CUG | 0.48 | 0.48 | 0.48 | 0.48 | 0.48 | 0.47 | 0.47 | 0.48 | 0.48 | 0.47 | 0.48 | 0.47 | 0.48 |
| Isoleucine(I) | AUU | 2.4 | 2.4 | 2.4 | 2.4 | 2.4 | 2.4 | 2.31 | 2.4 | 2.4 | 2.4 | 2.4 | 2.4 | 2.4 |
|  | AUC | 0 | 0 | 0 | 0 | 0 | 0 | 0.12 | 0 | 0 | 0 | 0 | 0 | 0 |
|  | AUA | 0.6 | 0.6 | 0.6 | 0.6 | 0.6 | 0.6 | 0.58 | 0.6 | 0.6 | 0.6 | 0.6 | 0.6 | 0.6 |
| Valine(V) | GUU | 1.49 | 1.49 | 1.49 | 1.49 | 1.49 | 1.55 | 1.43 | 1.49 | 1.49 | 1.55 | 1.49 | 1.55 | 1.49 |
|  | GUC | 0.09 | 0.09 | 0.09 | 0.09 | 0.09 | 0.09 | 0.1 | 0.09 | 0.09 | 0.09 | 0.09 | 0.09 | 0.09 |
|  | GUA | 1.21 | 1.21 | 1.21 | 1.21 | 1.21 | 1.18 | 1.24 | 1.12 | 1.21 | 1.18 | 1.21 | 1.18 | 1.21 |
|  | GUG | 1.21 | 1.21 | 1.21 | 1.21 | 1.21 | 1.18 | 1.24 | 1.3 | 1.21 | 1.18 | 1.21 | 1.18 | 1.21 |
| Serine(S) | UCU | 2.73 | 2.73 | 2.73 | 2.73 | 2.73 | 2.73 | 2.67 | 2.73 | 2.73 | 2.73 | 2.73 | 2.73 | 2.73 |
|  | UCC | 0.68 | 0.68 | 0.68 | 0.68 | 0.68 | 0.68 | 0.67 | 0.68 | 0.68 | 0.68 | 0.68 | 0.68 | 0.68 |
|  | UCA | 0.27 | 0.27 | 0.27 | 0.27 | 0.27 | 0.27 | 0.27 | 0.27 | 0.27 | 0.27 | 0.27 | 0.27 | 0.27 |
|  | UCG | 0.14 | 0.14 | 0.14 | 0.14 | 0.14 | 0.14 | 0.13 | 0.14 | 0.14 | 0.14 | 0.14 | 0.14 | 0.14 |
|  | AGU | 1.5 | 1.5 | 1.5 | 1.5 | 1.5 | 1.5 | 1.6 | 1.5 | 1.5 | 1.5 | 1.5 | 1.5 | 1.5 |
|  | AGC | 0.68 | 0.68 | 0.68 | 0.68 | 0.68 | 0.68 | 0.67 | 0.68 | 0.68 | 0.68 | 0.68 | 0.68 | 0.68 |
| Proline(P) | CCU | 2.09 | 2.09 | 2.09 | 2.04 | 2.09 | 2.09 | 2.14 | 2.04 | 2.09 | 2.09 | 2.09 | 2.04 | 2.09 |
|  | CCC | 0.73 | 0.73 | 0.73 | 0.8 | 0.73 | 0.73 | 0.65 | 0.8 | 0.73 | 0.73 | 0.73 | 0.71 | 0.73 |
|  | CCA | 1 | 1 | 1 | 0.98 | 1 | 1 | 1.12 | 0.98 | 1 | 1 | 1 | 1.07 | 1 |
|  | CCG | 0.18 | 0.18 | 0.18 | 0.18 | 0.18 | 0.18 | 0.09 | 0.18 | 0.18 | 0.18 | 0.18 | 0.18 | 0.18 |
| Threonine(T) | ACU | 1.78 | 1.78 | 1.89 | 1.67 | 1.78 | 1.78 | 1.84 | 1.78 | 1.78 | 1.78 | 1.71 | 1.78 | 1.78 |
|  | ACC | 0.89 | 0.89 | 0.89 | 0.89 | 0.89 | 0.89 | 0.86 | 0.89 | 0.89 | 0.89 | 0.91 | 0.89 | 0.89 |
|  | ACA | 1.11 | 1.11 | 1 | 1.11 | 1.11 | 1.11 | 1.08 | 1.11 | 1.11 | 1.11 | 1.14 | 1.11 | 1.11 |
|  | ACG | 0.22 | 0.22 | 0.22 | 0.33 | 0.22 | 0.22 | 0.22 | 0.22 | 0.22 | 0.22 | 0.23 | 0.22 | 0.22 |
| Alanine(A) | GCU | 1.94 | 1.94 | 1.94 | 1.94 | 1.94 | 1.87 | 2.06 | 1.94 | 1.94 | 1.87 | 2 | 1.87 | 1.94 |
|  | GCC | 1.03 | 1.03 | 1.03 | 1.03 | 1.03 | 1.07 | 0.9 | 1.03 | 1.03 | 1.07 | 1 | 1.07 | 1.03 |
|  | GCA | 1.03 | 1.03 | 1.03 | 1.03 | 1.03 | 1.07 | 1.03 | 1.03 | 1.03 | 1.07 | 1 | 1.07 | 1.03 |
|  | GCG | 0 | 0 | 0 | 0 | 0 | 0 | 0 | 0 | 0 | 0 | 0 | 0 | 0 |
| Tyrosine(Y) | UAU | 1.79 | 1.79 | 1.79 | 1.79 | 1.79 | 1.79 | 1.72 | 1.79 | 1.79 | 1.79 | 1.79 | 1.79 | 1.79 |
|  | UAC | 0.21 | 0.21 | 0.21 | 0.21 | 0.21 | 0.21 | 0.28 | 0.21 | 0.21 | 0.21 | 0.21 | 0.21 | 0.21 |
| Histidine(H) | CAU | 1.5 | 1.5 | 1.5 | 1.6 | 1.5 | 1.5 | 1.5 | 1.6 | 1.5 | 1.5 | 1.5 | 1.5 | 1.5 |
|  | CAC | 0.5 | 0.5 | 0.5 | 0.4 | 0.5 | 0.5 | 0.5 | 0.4 | 0.5 | 0.5 | 0.5 | 0.5 | 0.5 |
| Glutamine(Q) | CAA | 0.74 | 0.74 | 0.74 | 0.74 | 0.74 | 0.74 | 0.74 | 0.74 | 0.74 | 0.77 | 0.74 | 0.69 | 0.74 |
|  | CAG | 1.26 | 1.26 | 1.26 | 1.26 | 1.26 | 1.26 | 1.26 | 1.26 | 1.26 | 1.23 | 1.26 | 1.31 | 1.26 |
| Asparagine(N) | AAU | 1.57 | 1.57 | 1.57 | 1.57 | 1.57 | 1.57 | 1.64 | 1.57 | 1.57 | 1.57 | 1.57 | 1.57 | 1.57 |
|  | AAC | 0.43 | 0.43 | 0.43 | 0.43 | 0.43 | 0.43 | 0.36 | 0.43 | 0.43 | 0.43 | 0.43 | 0.43 | 0.43 |
| Lysine(K) | AAA | 0.96 | 0.96 | 0.96 | 0.96 | 0.96 | 0.96 | 1 | 0.96 | 0.96 | 0.96 | 0.96 | 0.96 | 0.96 |
|  | AAG | 1.04 | 1.04 | 1.04 | 1.04 | 1.04 | 1.04 | 1 | 1.04 | 1.04 | 1.04 | 1.04 | 1.04 | 1.04 |
| Aspartic acid(D) | GAU | 1.49 | 1.49 | 1.49 | 1.49 | 1.49 | 1.49 | 1.49 | 1.49 | 1.49 | 1.49 | 1.49 | 1.49 | 1.49 |
|  | GAC | 0.51 | 0.51 | 0.51 | 0.51 | 0.51 | 0.51 | 0.51 | 0.51 | 0.51 | 0.51 | 0.51 | 0.51 | 0.51 |
| Glutamic acid(E) | GAA | 1.29 | 1.29 | 1.29 | 1.29 | 1.29 | 1.29 | 1.43 | 1.29 | 1.29 | 1.29 | 1.29 | 1.29 | 1.29 |
|  | GAG | 0.71 | 0.71 | 0.71 | 0.71 | 0.71 | 0.71 | 0.57 | 0.71 | 0.71 | 0.71 | 0.71 | 0.71 | 0.71 |
| Cysteine C | UGU | 1.5 | 1.5 | 1.5 | 1.5 | 1.5 | 1.5 | 1.5 | 1.5 | 1.5 | 1.5 | 1.5 | 1.5 | 1.5 |
|  | UGC | 0.5 | 0.5 | 0.5 | 0.5 | 0.5 | 0.5 | 0.5 | 0.5 | 0.5 | 0.5 | 0.5 | 0.5 | 0.5 |
| Arginine R | CGU | 2.4 | 2.4 | 2.4 | 2.4 | 2.4 | 2.4 | 2.9 | 2.4 | 2.4 | 2.32 | 2.4 | 2.4 | 2.4 |
|  | CGC | 0.8 | 0.8 | 0.8 | 0.8 | 0.8 | 0.8 | 0.41 | 0.8 | 0.8 | 0.77 | 0.8 | 0.8 | 0.8 |
|  | CGA | 0 | 0 | 0 | 0 | 0 | 0 | 0 | 0 | 0 | 0 | 0 | 0 | 0 |
|  | CGG | 0.6 | 0.6 | 0.6 | 0.6 | 0.6 | 0.6 | 0.83 | 0.6 | 0.6 | 0.77 | 0.6 | 0.6 | 0.6 |
|  | AGA | 1 | 1 | 1 | 0.8 | 1 | 1.2 | 1.03 | 1 | 1 | 1.16 | 1 | 1 | 1 |
|  | AGG | 1.2 | 1.2 | 1.2 | 1.4 | 1.2 | 1 | 0.83 | 1.2 | 1.2 | 0.97 | 1.2 | 1.2 | 1.2 |
| Glycine (G) | GGU | 1.6 | 1.6 | 1.6 | 1.6 | 1.6 | 1.49 | 1.71 | 1.6 | 1.6 | 1.49 | 1.6 | 1.6 | 1.6 |
|  | GGC | 1.49 | 1.49 | 1.49 | 1.49 | 1.49 | 1.49 | 1.49 | 1.49 | 1.49 | 1.49 | 1.49 | 1.49 | 1.49 |
|  | GGA | 0.46 | 0.46 | 0.46 | 0.46 | 0.46 | 0.46 | 0.34 | 0.46 | 0.46 | 0.46 | 0.46 | 0.46 | 0.46 |
|  | GGG | 0.46 | 0.46 | 0.46 | 0.46 | 0.46 | 0.57 | 0.46 | 0.46 | 0.46 | 0.57 | 0.46 | 0.46 | 0.46 |

**Supplementary table 2.8:** Values of Relative Synonymous Codon Usage (RSCU) for accession number from KU707809.1 to KU707821.1 of L1 protein of Human Papillomavirus 18.

| **Amino acids** | **Codons** | **KU707809.1** | **KU707810.1** | **KU707811.1** | **KU707812.1** | **KU707813.1** | **KU707814.1** | **KU707815.1** | **KU707816.1** | **KU707817.1** | **KU707818.1** | **KU707819.1** | **KU707820.1** | **KU707821.1** |
| --- | --- | --- | --- | --- | --- | --- | --- | --- | --- | --- | --- | --- | --- | --- |
| Phenylalanine(F) | UUU | 1.83 | 1.83 | 1.83 | 1.83 | 1.83 | 1.83 | 1.83 | 1.75 | 1.75 | 1.83 | 1.75 | 1.83 | 1.83 |
|  | UUC | 0.17 | 0.17 | 0.17 | 0.17 | 0.17 | 0.17 | 0.17 | 0.25 | 0.25 | 0.17 | 0.25 | 0.17 | 0.17 |
| Leucine(L) | UUA | 2.82 | 2.88 | 2.82 | 2.82 | 2.94 | 2.88 | 2.82 | 2.82 | 2.82 | 2.82 | 2.82 | 2.94 | 2.82 |
|  | UUG | 1.65 | 1.56 | 1.65 | 1.65 | 1.53 | 1.56 | 1.65 | 1.59 | 1.59 | 1.65 | 1.59 | 1.53 | 1.65 |
|  | CUU | 0.59 | 0.6 | 0.59 | 0.59 | 0.59 | 0.6 | 0.59 | 0.61 | 0.61 | 0.59 | 0.61 | 0.59 | 0.59 |
|  | CUC | 0 | 0 | 0 | 0 | 0 | 0 | 0 | 0 | 0 | 0 | 0 | 0 | 0 |
|  | CUA | 0.47 | 0.48 | 0.47 | 0.47 | 0.47 | 0.48 | 0.47 | 0.49 | 0.49 | 0.47 | 0.49 | 0.47 | 0.47 |
|  | CUG | 0.47 | 0.48 | 0.47 | 0.47 | 0.47 | 0.48 | 0.47 | 0.49 | 0.49 | 0.47 | 0.49 | 0.47 | 0.47 |
| Isoleucine(I) | AUU | 2.4 | 2.4 | 2.4 | 2.4 | 2.31 | 2.4 | 2.4 | 2.4 | 2.4 | 2.4 | 2.4 | 2.4 | 2.4 |
|  | AUC | 0 | 0 | 0 | 0 | 0.12 | 0 | 0 | 0 | 0 | 0 | 0 | 0 | 0 |
|  | AUA | 0.6 | 0.6 | 0.6 | 0.6 | 0.58 | 0.6 | 0.6 | 0.6 | 0.6 | 0.6 | 0.6 | 0.6 | 0.6 |
| Valine(V) | GUU | 1.55 | 1.49 | 1.49 | 1.49 | 1.43 | 1.49 | 1.45 | 1.49 | 1.49 | 1.55 | 1.49 | 1.55 | 1.49 |
|  | GUC | 0.09 | 0.09 | 0.09 | 0.09 | 0.1 | 0.09 | 0.09 | 0.09 | 0.09 | 0.09 | 0.09 | 0.09 | 0.09 |
|  | GUA | 1.18 | 1.21 | 1.21 | 1.21 | 1.24 | 1.21 | 1.18 | 1.21 | 1.21 | 1.18 | 1.21 | 1.18 | 1.21 |
|  | GUG | 1.18 | 1.21 | 1.21 | 1.21 | 1.24 | 1.21 | 1.27 | 1.21 | 1.21 | 1.18 | 1.21 | 1.18 | 1.21 |
| Serine(S) | UCU | 2.73 | 2.73 | 2.73 | 2.73 | 2.67 | 2.73 | 2.73 | 2.73 | 2.73 | 2.73 | 2.73 | 2.73 | 2.73 |
|  | UCC | 0.68 | 0.68 | 0.68 | 0.68 | 0.67 | 0.68 | 0.68 | 0.68 | 0.68 | 0.68 | 0.68 | 0.68 | 0.68 |
|  | UCA | 0.27 | 0.27 | 0.27 | 0.27 | 0.27 | 0.27 | 0.27 | 0.27 | 0.27 | 0.27 | 0.27 | 0.27 | 0.27 |
|  | UCG | 0.14 | 0.14 | 0.14 | 0.14 | 0.13 | 0.14 | 0.14 | 0.14 | 0.14 | 0.14 | 0.14 | 0.14 | 0.14 |
|  | AGU | 1.5 | 1.5 | 1.5 | 1.5 | 1.6 | 1.5 | 1.5 | 1.5 | 1.5 | 1.5 | 1.5 | 1.5 | 1.5 |
|  | AGC | 0.68 | 0.68 | 0.68 | 0.68 | 0.67 | 0.68 | 0.68 | 0.68 | 0.68 | 0.68 | 0.68 | 0.68 | 0.68 |
| Proline(P) | CCU | 2.04 | 2.09 | 2.09 | 2.04 | 2.14 | 2.09 | 2.04 | 2.09 | 2.09 | 2.04 | 2.09 | 2.04 | 2.09 |
|  | CCC | 0.71 | 0.73 | 0.73 | 0.71 | 0.65 | 0.73 | 0.71 | 0.73 | 0.73 | 0.71 | 0.73 | 0.71 | 0.73 |
|  | CCA | 1.07 | 1 | 1 | 0.98 | 1.12 | 1 | 1.07 | 1 | 1 | 1.07 | 1 | 1.07 | 1 |
|  | CCG | 0.18 | 0.18 | 0.18 | 0.27 | 0.09 | 0.18 | 0.18 | 0.18 | 0.18 | 0.18 | 0.18 | 0.18 | 0.18 |
| Threonine(T) | ACU | 1.67 | 1.78 | 1.78 | 1.78 | 1.84 | 1.78 | 1.78 | 1.78 | 1.78 | 1.67 | 1.78 | 1.78 | 1.78 |
|  | ACC | 1 | 0.89 | 0.89 | 0.89 | 0.86 | 0.89 | 0.89 | 0.89 | 0.89 | 1 | 0.89 | 0.89 | 0.89 |
|  | ACA | 1.11 | 1.11 | 1.11 | 1.11 | 1.08 | 1.11 | 1.11 | 1.11 | 1.11 | 1.11 | 1.11 | 1.11 | 1.11 |
|  | ACG | 0.22 | 0.22 | 0.22 | 0.22 | 0.22 | 0.22 | 0.22 | 0.22 | 0.22 | 0.22 | 0.22 | 0.22 | 0.22 |
| Alanine(A) | GCU | 1.87 | 1.94 | 1.94 | 1.94 | 2.06 | 1.94 | 1.87 | 1.94 | 1.94 | 1.87 | 1.94 | 1.87 | 1.94 |
|  | GCC | 1.07 | 1.03 | 1.03 | 1.03 | 0.9 | 1.03 | 1.07 | 1.03 | 1.03 | 1.07 | 1.03 | 1.07 | 1.03 |
|  | GCA | 1.07 | 1.03 | 1.03 | 1.03 | 1.03 | 1.03 | 1.07 | 1.03 | 1.03 | 1.07 | 1.03 | 1.07 | 1.03 |
|  | GCG | 0 | 0 | 0 | 0 | 0 | 0 | 0 | 0 | 0 | 0 | 0 | 0 | 0 |
| Tyrosine(Y) | UAU | 1.79 | 1.79 | 1.79 | 1.79 | 1.72 | 1.79 | 1.79 | 1.79 | 1.79 | 1.79 | 1.79 | 1.79 | 1.79 |
|  | UAC | 0.21 | 0.21 | 0.21 | 0.21 | 0.28 | 0.21 | 0.21 | 0.21 | 0.21 | 0.21 | 0.21 | 0.21 | 0.21 |
| Histidine(H) | CAU | 1.5 | 1.5 | 1.5 | 1.5 | 1.5 | 1.5 | 1.5 | 1.5 | 1.5 | 1.5 | 1.5 | 1.5 | 1.5 |
|  | CAC | 0.5 | 0.5 | 0.5 | 0.5 | 0.5 | 0.5 | 0.5 | 0.5 | 0.5 | 0.5 | 0.5 | 0.5 | 0.5 |
| Glutamine(Q) | CAA | 0.69 | 0.74 | 0.74 | 0.69 | 0.74 | 0.74 | 0.69 | 0.74 | 0.74 | 0.69 | 0.74 | 0.69 | 0.74 |
|  | CAG | 1.31 | 1.26 | 1.26 | 1.31 | 1.26 | 1.26 | 1.31 | 1.26 | 1.26 | 1.31 | 1.26 | 1.31 | 1.26 |
| Asparagine(N) | AAU | 1.57 | 1.57 | 1.57 | 1.57 | 1.64 | 1.57 | 1.57 | 1.57 | 1.57 | 1.57 | 1.57 | 1.57 | 1.57 |
|  | AAC | 0.43 | 0.43 | 0.43 | 0.43 | 0.36 | 0.43 | 0.43 | 0.43 | 0.43 | 0.43 | 0.43 | 0.43 | 0.43 |
| Lysine(K) | AAA | 0.96 | 0.96 | 0.96 | 0.96 | 1 | 0.96 | 0.96 | 0.96 | 0.96 | 0.96 | 0.96 | 0.96 | 1 |
|  | AAG | 1.04 | 1.04 | 1.04 | 1.04 | 1 | 1.04 | 1.04 | 1.04 | 1.04 | 1.04 | 1.04 | 1.04 | 1 |
| Aspartic acid(D) | GAU | 1.49 | 1.49 | 1.49 | 1.49 | 1.49 | 1.49 | 1.49 | 1.49 | 1.49 | 1.49 | 1.49 | 1.49 | 1.49 |
|  | GAC | 0.51 | 0.51 | 0.51 | 0.51 | 0.51 | 0.51 | 0.51 | 0.51 | 0.51 | 0.51 | 0.51 | 0.51 | 0.51 |
| Glutamic acid(E) | GAA | 1.29 | 1.29 | 1.29 | 1.29 | 1.43 | 1.29 | 1.29 | 1.29 | 1.29 | 1.29 | 1.29 | 1.29 | 1.29 |
|  | GAG | 0.71 | 0.71 | 0.71 | 0.71 | 0.57 | 0.71 | 0.71 | 0.71 | 0.71 | 0.71 | 0.71 | 0.71 | 0.71 |
| Cysteine C | UGU | 1.5 | 1.5 | 1.5 | 1.5 | 1.5 | 1.5 | 1.5 | 1.5 | 1.5 | 1.5 | 1.5 | 1.5 | 1.5 |
|  | UGC | 0.5 | 0.5 | 0.5 | 0.5 | 0.5 | 0.5 | 0.5 | 0.5 | 0.5 | 0.5 | 0.5 | 0.5 | 0.5 |
| Arginine R | CGU | 2.4 | 2.4 | 2.4 | 2.4 | 2.9 | 2.4 | 2.4 | 2.4 | 2.4 | 2.4 | 2.4 | 2.4 | 2.32 |
|  | CGC | 0.8 | 0.8 | 0.8 | 0.8 | 0.41 | 0.8 | 0.8 | 0.8 | 0.8 | 0.8 | 0.8 | 0.8 | 0.77 |
|  | CGA | 0 | 0 | 0 | 0 | 0 | 0 | 0 | 0 | 0 | 0 | 0 | 0 | 0 |
|  | CGG | 0.6 | 0.6 | 0.6 | 0.6 | 0.83 | 0.6 | 0.6 | 0.6 | 0.6 | 0.6 | 0.6 | 0.6 | 0.58 |
|  | AGA | 1 | 1 | 1 | 1 | 1.03 | 1 | 1 | 1 | 1 | 1 | 1 | 1 | 0.97 |
|  | AGG | 1.2 | 1.2 | 1.2 | 1.2 | 0.83 | 1.2 | 1.2 | 1.2 | 1.2 | 1.2 | 1.2 | 1.2 | 1.35 |
| Glycine (G) | GGU | 1.6 | 1.6 | 1.6 | 1.6 | 1.71 | 1.6 | 1.6 | 1.6 | 1.6 | 1.6 | 1.6 | 1.6 | 1.6 |
|  | GGC | 1.49 | 1.49 | 1.49 | 1.49 | 1.49 | 1.49 | 1.49 | 1.49 | 1.49 | 1.49 | 1.49 | 1.49 | 1.49 |
|  | GGA | 0.46 | 0.46 | 0.46 | 0.46 | 0.34 | 0.46 | 0.46 | 0.46 | 0.46 | 0.46 | 0.46 | 0.46 | 0.46 |
|  | GGG | 0.46 | 0.46 | 0.46 | 0.46 | 0.46 | 0.46 | 0.46 | 0.46 | 0.46 | 0.46 | 0.46 | 0.46 | 0.46 |

**Supplementary table 2.9:** Values of Relative Synonymous Codon Usage (RSCU) for accession number from KU707822.1 to KU707825.1 of L1 protein of Human Papillomavirus 18.

| **Amino acids** | **Codons** | **KU707822.1** | **KU707823.1** | **KU707824.1** | **KU707825.1** |
| --- | --- | --- | --- | --- | --- |
| Phenylalanine(F) | UUU | 1.83 | 1.83 | 1.83 | 1.83 |
|  | UUC | 0.17 | 0.17 | 0.17 | 0.17 |
| Leucine(L) | UUA | 2.82 | 2.88 | 2.88 | 2.77 |
|  | UUG | 1.65 | 1.56 | 1.56 | 1.62 |
|  | CUU | 0.59 | 0.6 | 0.6 | 0.69 |
|  | CUC | 0 | 0 | 0 | 0 |
|  | CUA | 0.47 | 0.48 | 0.48 | 0.46 |
|  | CUG | 0.47 | 0.48 | 0.48 | 0.46 |
| Isoleucine(I) | AUU | 2.4 | 2.4 | 2.4 | 2.38 |
|  | AUC | 0 | 0 | 0 | 0 |
|  | AUA | 0.6 | 0.6 | 0.6 | 0.62 |
| Valine(V) | GUU | 1.49 | 1.49 | 1.49 | 1.49 |
|  | GUC | 0.09 | 0.09 | 0.09 | 0.09 |
|  | GUA | 1.21 | 1.21 | 1.21 | 1.21 |
|  | GUG | 1.21 | 1.21 | 1.21 | 1.21 |
| Serine(S) | UCU | 2.73 | 2.73 | 2.73 | 2.73 |
|  | UCC | 0.68 | 0.68 | 0.68 | 0.68 |
|  | UCA | 0.27 | 0.27 | 0.27 | 0.27 |
|  | UCG | 0.14 | 0.14 | 0.14 | 0.14 |
|  | AGU | 1.5 | 1.5 | 1.5 | 1.5 |
|  | AGC | 0.68 | 0.68 | 0.68 | 0.68 |
| Proline(P) | CCU | 2.09 | 2.09 | 2.09 | 2.09 |
|  | CCC | 0.73 | 0.73 | 0.73 | 0.73 |
|  | CCA | 1 | 1 | 1 | 1 |
|  | CCG | 0.18 | 0.18 | 0.18 | 0.18 |
| Threonine(T) | ACU | 1.78 | 1.78 | 1.78 | 1.78 |
|  | ACC | 0.89 | 0.89 | 0.89 | 0.89 |
|  | ACA | 1.11 | 1.11 | 1.11 | 1.11 |
|  | ACG | 0.22 | 0.22 | 0.22 | 0.22 |
| Alanine(A) | GCU | 1.94 | 1.94 | 1.94 | 1.94 |
|  | GCC | 1.03 | 1.03 | 1.03 | 1.03 |
|  | GCA | 1.03 | 1.03 | 1.03 | 1.03 |
|  | GCG | 0 | 0 | 0 | 0 |
| Tyrosine(Y) | UAU | 1.79 | 1.79 | 1.79 | 1.79 |
|  | UAC | 0.21 | 0.21 | 0.21 | 0.21 |
| Histidine(H) | CAU | 1.5 | 1.5 | 1.47 | 1.5 |
|  | CAC | 0.5 | 0.5 | 0.53 | 0.5 |
| Glutamine(Q) | CAA | 0.74 | 0.74 | 0.74 | 0.74 |
|  | CAG | 1.26 | 1.26 | 1.26 | 1.26 |
| Asparagine(N) | AAU | 1.57 | 1.57 | 1.57 | 1.57 |
|  | AAC | 0.43 | 0.43 | 0.43 | 0.43 |
| Lysine(K) | AAA | 1 | 0.96 | 0.96 | 0.96 |
|  | AAG | 1 | 1.04 | 1.04 | 1.04 |
| Aspartic acid(D) | GAU | 1.49 | 1.49 | 1.49 | 1.49 |
|  | GAC | 0.51 | 0.51 | 0.51 | 0.51 |
| Glutamic acid(E) | GAA | 1.29 | 1.29 | 1.29 | 1.29 |
|  | GAG | 0.71 | 0.71 | 0.71 | 0.71 |
| Cysteine C | UGU | 1.5 | 1.5 | 1.5 | 1.5 |
|  | UGC | 0.5 | 0.5 | 0.5 | 0.5 |
| Arginine R | CGU | 2.32 | 2.4 | 2.52 | 2.4 |
|  | CGC | 0.77 | 0.8 | 0.77 | 0.8 |
|  | CGA | 0 | 0 | 0 | 0 |
|  | CGG | 0.58 | 0.6 | 0.58 | 0.8 |
|  | AGA | 0.97 | 1 | 0.97 | 1 |
|  | AGG | 1.35 | 1.2 | 1.16 | 1 |
| Glycine (G) | GGU | 1.6 | 1.6 | 1.6 | 1.6 |
|  | GGC | 1.49 | 1.49 | 1.49 | 1.49 |
|  | GGA | 0.46 | 0.46 | 0.46 | 0.46 |
|  | GGG | 0.46 | 0.46 | 0.46 | 0.46 |
